# A transient third metal ion shapes the catalytic cycle of a group I intron ribozyme

**DOI:** 10.64898/2026.10.07.757386

**Authors:** Erika McCarthy, Şölen Ekesan, Timothy J. Giese, Yanan Du, German P. Barletta, Andreas W. Götz, Joseph A. Piccirilli, Darrin M. York

**Affiliations:** Laboratory for Biomolecular Simulation Research, Institute for Quantitative Biomedicine and Department of Chemistry and Chemical Biology, Rutgers University, Piscataway, New Jersey 08854, United States; San Diego Supercomputer Center, University of California San Diego, La Jolla, California 92093-0505, United States; Department of Chemistry and Department of Biochemistry and Molecular Biology, The University of Chicago, Chicago, Illinois 60637, United States

## Abstract

The self-splicing group I intron ribozyme from *Tetrahymena thermophillia* (THr) is a prototypical example of how RNA overcomes its limited chemical repertoire to form complex, negatively charged junctions and catalyze chemical reactions. Despite over 30 years of biochemical and structural characterization, questions still remain regarding the requirements for THr catalysis that may inform structure based drug design against group I introns, as well as the use of THr as an RNA editing tool. Recent cryo-EM structures have provided an unprecedented window into the first and second steps of splicing, enabling detailed mechanistic investigation at the atomistic level. In this work, we leverage solvation theory, molecular dynamics, alchemical free energy simulations, and machine learning accelerated QM/MM to probe outstanding questions related to 1) the role of a possible third Mg^2+^ binding in the active site, 2) differences in catalytic requirements between G_exo_ cofactors and between the first and second steps of splicing, 3) the potential role of an acid or base, and 4) restrictions on the identity of the 5*^′^* exon splice site nucleotide, U(−1). We propose that a transient, cofactor dependent, interaction with a third metal participates in the first step, while a higher affinity interaction with this third ion is required in the second step. We also propose that the presence of a phosphate on the G_exo_ in the first step, and analogously the G*_ω_* in the second step, forms a metal ion binding site not present when guanosine is bound. Finally, we propose and computationally test design modifications of G22 (isofunctional substitutions of 2,6-diaminopurine that tune the p*K*_a_ at the N1 position) that we predict will enable the accommodation of C(–1) in a crucial wobble-like pair at physiological pH. This design would provide a potential strategy for expanding the sequence space accessible to programmable trans-splicing ribozymes in RNA editing applications.

**TOC Graphic:** 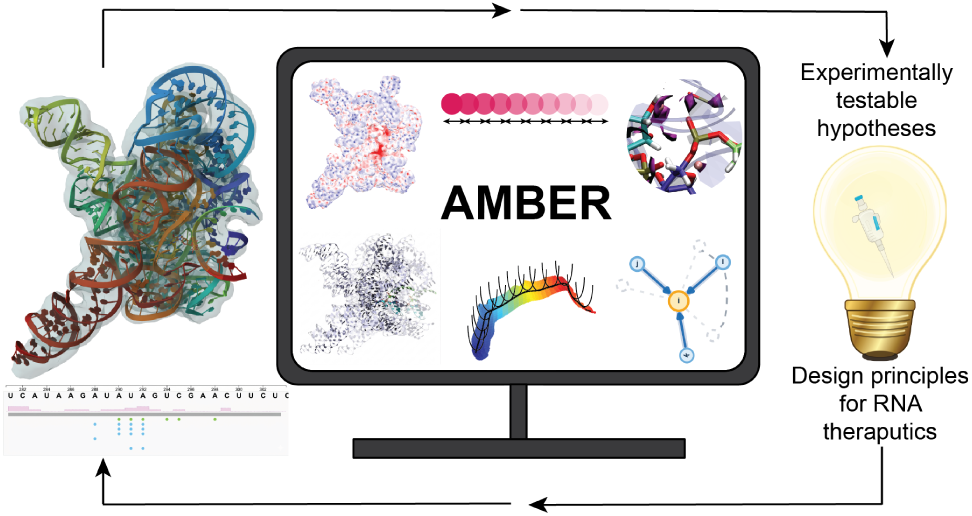

## Introduction

Group I introns are non-coding RNA enzymes (ribozymes) that catalyze their own excision and subsequent ligation of neighboring exons through two sequential transesterification reactions, a process known as splicing.^1^ These ribozymes are vitally important as a foundational model for RNA catalysis and self-splicing that underpins modern biotechnology tools and informs RNA based therapeutic strategies in medicine.^2–5^ The discovery of self-splicing activity by the group I intron ribozyme from *Tetrahymena thermophila* (THr) in 1982 was the first direct evidence that RNA can act as an enzyme in the absence of protein cofactors.^1,6,7^ This finding marked a paradigm shift in RNA’s perceived role in the central dogma of molecular biology and opened entirely new fields of inquiry into RNA structure, mechanism, and evolution.^8,9^

Since this landmark discovery, the known landscape of RNA chemistry has expanded dramatically.^10–13^ Naturally occurring ribozymes that catalyze phosphoryl transfer have been discovered across eukaryotes,^7,14–17^ viruses,^18,19^ fungi,^20,21^ and bacteria.^22–25^ More diverse chemistry has been artificially evolved in synthetic nucleic acid enzymes capable of catalyzing phosphoryl transfer,^26,27^ alkyl transfer,^28–33^ Diels–Alder reaction,^34,35^ Michael addition,^36^ and aldol condensation.^37^

Across these systems, electrostatics are key for catalysis,^38^ often governing the recruitment and positioning of divalent metal ions that are essential for both structural integrity and catalytic function.^10^ For small RNA-cleaving nucleic acid enzymes, shared mechanistic strategies^39–41^ and conserved active site motifs^38,42,43^ have begun to emerge, establishing a conceptual framework to inform and guide rational design. For more complex catalytic RNA systems, such as group I introns, a deeper mechanistic understanding is desired to solidify the catalytic picture in atomic detail. The goal of the current study is to provide new mechanistic insight into THr as an important archetype group I intron model system, building upon a wealth of experimental data, including recent high-resolution cryo-EM structures.^44–46^

The L-16 S*ca*I ribozyme is a linear construct of THr capable of catalyzing the cleavage and ligation reaction in trans by binding an exogenous guanine cofactor, termed G_exo_, and a substrate oligonucleotide, bound by the internal guide sequence (IGS) of the ribozyme.^47^ An overview of the splicing reaction is given in Figure 1a. The first step of the reaction involves the cleavage of the O3*^′^*-P bond connecting the 3*^′^* end of the 5*^′^* exon and the 5*^′^* end of the intron and requires the binding of G_exo_, which contains the nucleophilic O3*^′^* (Figure 1b). The second step of the reaction involves the ligation of the 5*^′^* and 3*^′^* exons and requires the cleavage of the O3*^′^*-P bond that connects the 3*^′^* end of the intron to the 5*^′^* end of the 3*^′^* exon (Figure 1c). This step requires the release of G_exo_ and binding of the G(−1) nucleotide of the intron, termed G*_ω_*, to the G binding site on the P7 helix. Such guanine recognition motifs play a central role across RNA classes from ribozyme cofactor binding to metabolite sensing by riboswitches.^29,48,49^ Recent cryo-EM data provide a high-resolution picture of the active site architecture along the cleavage^46^ and ligation^44,45^ pathways (Figure 1d,e). This data builds on pioneering X-ray crystal structures^50–52^ by resolving peripheral domains that stabilize the core through tertiary interactions.^53,54^ These new structures provide an unprecedented window into THr catalysis, enabling mechanistic investigation at the atomic level using advanced computational methods. A schematic overview of both cis and trans-splicing as well as the ribozyme domains are given in Figures S1 and S2 of the Supporting Information.

**Figure 1.**
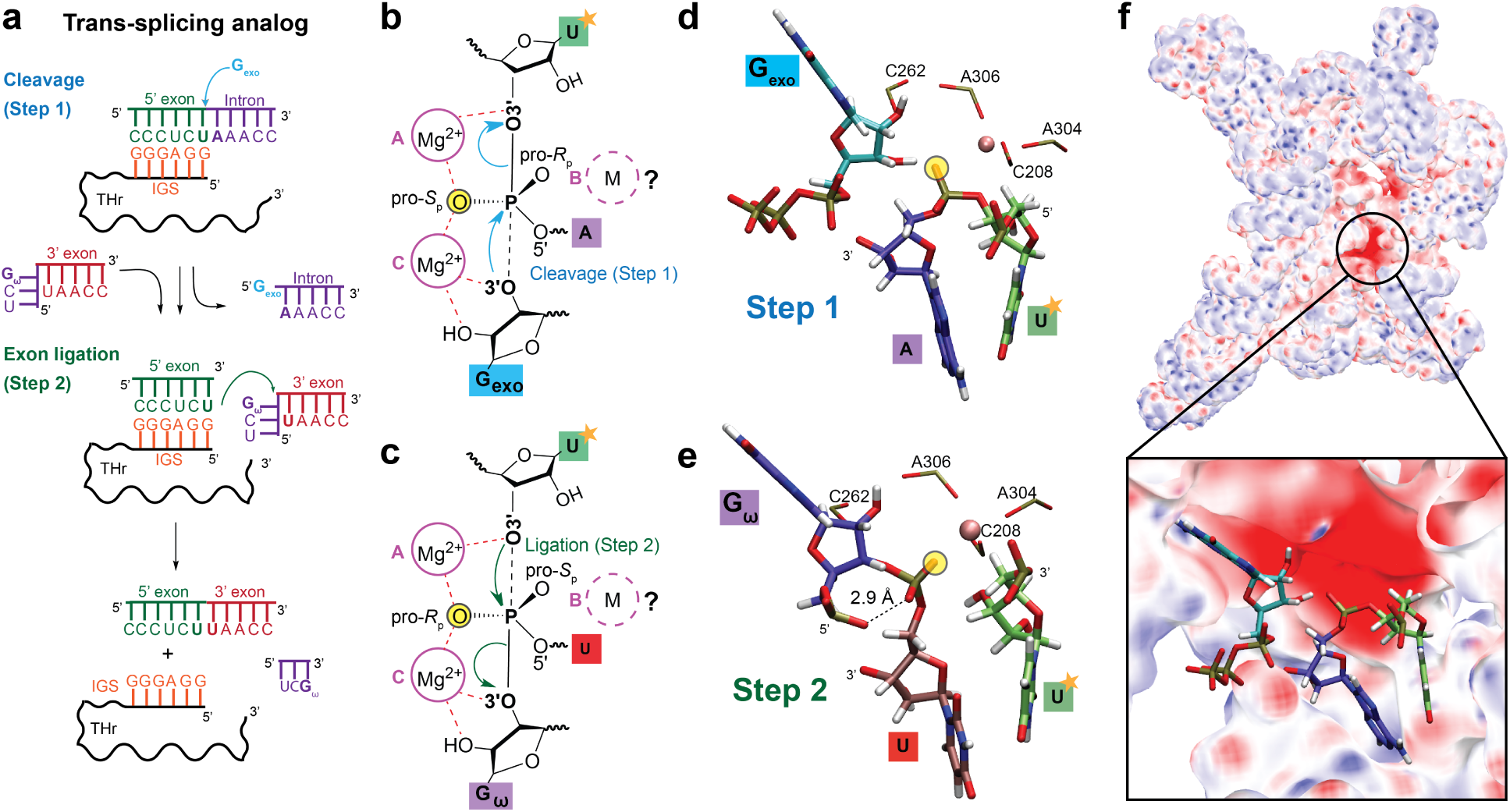
Overview of the THr catalyzed trans-splicing reaction. **(a)** Schematic of the transsplicing reaction analog catalyzed by THr from which cryo-EM structures where captured. Schematics of the **(b)** cleavage step (Step 1) and **(c)** ligation step (Step 2) phosphoryl transfer reactions with labeled metal ion binding sites. A yellow circle indicates a thio substitution present from the cryo-EM structure. Cryo-EM resolved active site of the **(d)** cleavage (7YCH) and **(e)** ligation (7YGA) steps of splicing, with active site phosphates (PO*^−^*) represented as sticks with residue labels. The phosphorothioate substitution used experimentally to slow the reaction is highlighted yellow. **(f)** Electrostatic potential surface representation of THr from the cleavage reaction from a Poisson-Boltzmann Surface Area calculation with the active site highlighted by a black box. Red coloring indicates negative charge, and blue positive charge.

However, the unambiguous identification of catalytically critical metal ion binding modes in ribozymes remains highly challenging.^55,56^ The active state is often transient and challenging to capture experimentally,^57,58^ or it is deliberately suppressed through chemical modification or altered experimental conditions to arrest activity and enable the conformational trapping needed for high-resolution structural determination. Moreover, catalytically relevant metal ion binding modes may be fractionally occupied and difficult to resolve, particularly for Mg^2+^ and Na^+^ ions, which are isoelectronic with water molecules; these ions may also be highly dynamic and rearrange or appear transiently along the reaction pathway.^59–61^

The THr active site is located within the ribozyme’s central P3–P7 catalytic core, where conserved helices and junctions converge to create an electrostatically repulsive pocket^38^ (Figure 1f) that binds the P1 substrate duplex and recruits catalytically critical Mg^2+^ ions. The exact coordination environments of active site Mg^2+^ ions (Figure 1b,c) in the cryo-EM structures, as well as the positions of key atoms of the scissile phosphate, are obscured due to a phosphorothioate substitution (Figure 1d,e), designed to prevent Mg^2+^ binding and inhibit activity, as is required for structural determination.^44–46^ The presence of one active site metal ion, Mg(A), was previously inferred from the THr crystal structure of Guo et al.^52^ based on a heavy metal soak. In the *Azoarcus* group I intron, which has a structurally similar active site scaffold to THr and similar guanosine binding kinetics,^62^ Mg(A) interaction with the leaving group O3*^′^* was inferred from thiophillic rescue experiments,^63^ and Mg(A) and Mg(C) positions were first resolved by Stahley et al.^64^ These have been further confirmed by recently by cryo-EM, which also resolved two conserved Mg^2+^ ions peripheral to the active site.^65^

Biochemical experiments on THr support at least two metal ions binding at the scissile phosphate. Phosphorothioate and metal ion rescue experiments have identified metal ion contacts in the cleavage reaction with the O3*^′^* nucleophile,^66^ leaving group,^67,68^ and pro-*S_P_*oxygen of the scissile phosphate (stereochemistry corresponding to the cleavage reaction).^69–71^ Additionally, the phosphates of A207 (pro-*S_P_*), C208 (pro-*S_P_*), C262 (pro-*S_P_*), A304 (pro-*S_P_*), and A306 (pro-*R_P_* and pro-*S_P_*) that form the active site make functionally important metal ion contacts.^62,64,72^ Double thio substitution/metal rescue experiments at the O3*^′^* of the substrate leaving group and pro-*S_P_* of the scissile phosphate in the cleavage reaction suggest these positions coordinate the same functionally important divalent metal ion,^73^ and amino substitution at the O2*^′^* position of G_exo_ suggests it also coordinates a metal.^74,75^ Whether the O3*^′^* nucleophile and O2*^′^* of G_exo_ in the cleavage reaction coordinate the same or distinct metal ions, implying 2 or 3 active site ions in total, has been a subject of considerable interest and debate.^66,71,76^ Traditionally, the metal ions coordinating the O3*^′^* of U(−1), O3*^′^* of G, and O2*^′^* of G were designated A, B, and C, respectively based on their thermodynamic fingerprints.^66,75^ However, later work showed that individual amino substitutions at the 3*^′^* and 2*^′^* positions of G yielded indistinguishable Mn^2+^ association constants and rescue effects, consistent with the thermodynamic fingerprint for Mg(C).^66,74,76^ Herein, the A and C designations will be maintained as outlined in ref 76, with B used more generally to refer to any additional active site binding mode not already biochemically characterized, for which first and second sphere coordination modes will be explored. Two Mg^2+^ ions were also resolved peripheral to the active site: Mg(D) which coordinates C262(pro-*R_P_*) and A207(pro-*S_P_*), and Mg(E) which coordinates U307(pro-*R_P_*) and A308(pro-*R_P_*). The Mg(D) nomenclature in this work is distinct from that of ref 77 and corresponds to M_2_ in ref 44, which is not equivalent to the M_2_ position in the *Azoarcus* ribozyme outlined in ref 78 (see Table S1 of the Supporting Information).

Finally, substitution at the pro-*R_P_*position in the cleavage reaction with guanosine bound yields a modest thio effect,^69,79^ which diminishes above pH 6.9 where a change in rate limiting step likely occurs,^80^ and this thio effect has not been measured with other G_exo_ moieties or for the pro-*S_P_* in the ligation step.^69^ The proximity of the pro-*R_P_* to the O3*^′^* leaving group and nucleophile in both cleavage and ligation, O2*^′^* of G_exo_, and lack of a rescue experiment begs the question as to whether it could interact with the elusive third metal ion (B) as depicted in Figure 1b,c. Three divalent metal ions have been implicated in the reactions of DNA polymerase *η* ^60^ and Ribonuclease P,^81,82^ and a dynamic interplay between divalent and monovalent ions has been highlighted in reactions of the RNase mitochondrial RNA processing (MRP) complex,^83^ spliceosome,^84,85^ group II intron,^86^ and RNase H1.^87^ The famous CRISPR-Cas9 system uses a two-Mg^2+^ mechanism for DNA cleavage, meanwhile a lysine residue stabilizes accumulated negative charge at the scissile phosphate.^88^

In this work, we apply an integrated computational enzymology approach that leverages recent advances in machine learning corrected quantum mechanical/molecular mechanical (QM/MM+ΔMLP) simulations,^89^ along with free energy surface^90^ and minimum free energy path^91^ sampling methods, alchemical free energy simulations,^92,93^ and molecular solvation theory^38^ to gain a predictive understanding of the THr mechanism. Our results shed light on the identity of acid and base catalysts for the cleavage and ligation steps of the reaction, rationalizing differences in catalytic requirements between G_exo_ cofactors and between the first and second step of splicing. Further, we provide evidence for the involvement of a third metal ion in catalysis that imparts electrostatic stabilization and solvent organization around the scissile phosphate. Finally, our calculations predict an isofunctional nucleobase substitution of the conserved G·U(−1) wobble pair between the 5*^′^* exon and IGS that would expand the range of splicing target sequences at neutral pH and enhance THr as a tool for sequence specific trans-splicing in therapeutic applications.

## Results and Discussion

We begin by analyzing electrostatic interactions in the active site from cryo-EM structures and Poisson-Boltzmann^94–96^ calculations, and identifying functional divalent (Mg^2+^) metal ion binding modes using molecular solvation theory (3D-RISM) and alchemical free energy (AFE) simulations. Next we use long timescale classical molecular dynamics (MD) simulations to ascertain the implications of these Mg^2+^ binding modes on the catalytic fitness and to characterize the active state dynamical ensemble in solution. Then, departing from the active state, we investigate the chemical steps of the reaction using Δ-machine learning accelerated quantum mechanical/molecular mechanical (QM/MM+ΔMLP) simulations to identify the minimum free energy pathways and transition state ensembles for different mechanisms. Finally, we perform computational mutagenesis simulations of an isofunctional substitution of the conserved G·U(−1) wobble in the cleavage reaction to enable predictions that have implications for biotechnology and therapeutic applications.

### An electronegative THr active site recruits structural and catalytic Mg^2+^ ions

The cryo-EM structures of Zhang,^46^ Li,^45^ and Su et al^44^ contain 3, 8, and 31 resolved Mg^2+^ ions with resolutions of 3.09, 2.35, and 3.05 Å, respectively. The structure of Zhang et al^46^ (PDB ID: 7YCH) represents the pre-catalytic state of the cleavage reaction (Step 1 of splicing), and those of Li et al^45^ (PDB ID: 7YGA) and Su et al^44^ (PDB ID: 7EZ2) represent the pre-catalytic state of the ligation reaction (Step 2 of splicing). The root mean square deviation between the backbone atoms of the ribozyme structures ranged between 1.72-2.46 Å. This high degree of structural overlap, especially in the core of the enzyme where metal ion binding is extremely important (see Figure S2 of the Supporting Information), enabled the mapping of important divalent metal ion binding sites between the structures. 7YCH served as a departure point for simulations of the cleavage reaction, and 7YGA, having the better resolution than 7EZ2, was chosen as the departure point for ligation step simulations. However, additional Mg^2+^ ions resolved in 7EZ2 were aligned onto 7YCH and 7YGA to preserve these important electrostatic interactions. The scissile phosphate oxygens of 7YCH and 7YGA only coordinate 1 Mg^2+^ ion, termed Mg(A) due to the de-activating thio substitution (highlighted in yellow in Figure 1d,e). Mg(C) was resolved in 7EZ2 and modeled in.

As shown in Figure 1f and Figure S3, the splice site resides within a junction of densely packed phosphate groups that generates a pronounced negative electrostatic potential, rendering site specific metal ion binding essential for assembling a catalytically competent active site and stabilizing the transition state. In the Step 2 structure (Figure 1e), the pro-*S_P_* oxygen of the scissile phosphate and the pro-*R_P_* oxygen of G*_ω_* approach within 2.9 Å, raising the possibility that a third metal ion mediates this contact via site B (Figure 1c). We thus set out to explore this hypothesis in the next two sections using molecular solvation theory and alchemical free energy simulations.

### 3D-RISM suggests a potential third Mg^2+^ binding site

The application of statistical mechanical molecular solvation theory,^97,98^ and in particular 3D-RISM,^99,100^ has provided key insight into metal ion binding in nucleic acids.^38,101–104^ We therefore applied 3D-RISM to the Step 1 and Step 2 pre-catalytic structures, stripped of solvent, to validate cryo-EM Mg^2+^ positions and probe for unresolved binding sites. In these calculations, thio substitutions in the cryo-EM structures were replaced by oxygen and the nucleophile was maintained in its neutral, standard state (SS). The calculations provide a ratio of the probability density of Mg^2+^ ion binding at a given site relative to bulk solution (*ρ*_site_/*ρ*_bulk_). The resulting ion distributions recapitulate sites A and C with high affinity in both steps (Figure 2a,b), confirming that these sites are intrinsic to the electrostatic architecture of the active site.

**Figure 2.**
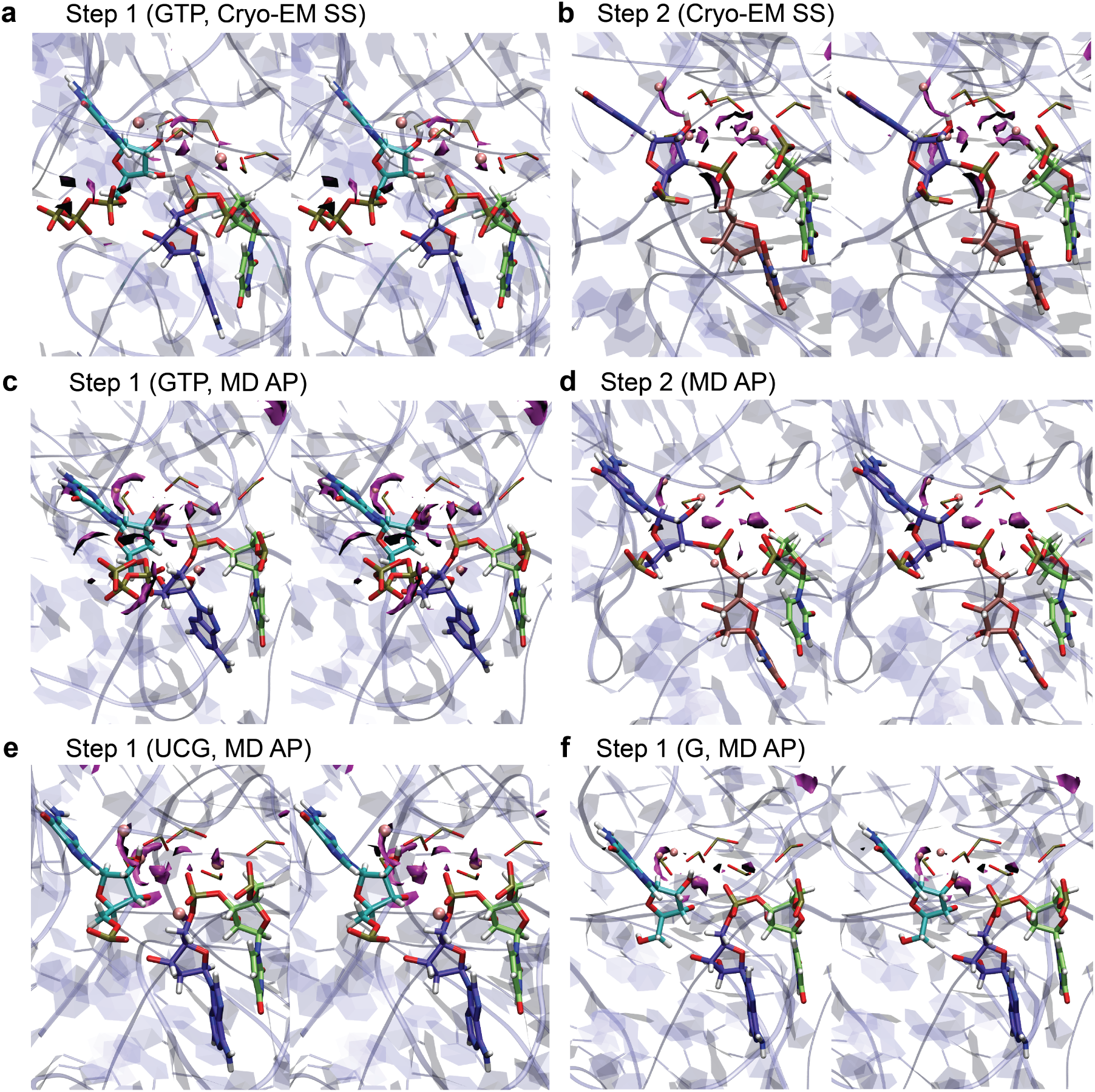
Active site Mg^2+^ ion isodensity contours predicted by 3D-RISM for **(a)** Step 1 (cleavage) and **(b)** Step 2 (ligation) in the standard state (SS) from cryo-EM with nucleophile protontated. Ions resolved in both cryo-EM structures are represented as spheres. 3D-RISM isodensity contours for **(c)** Step 1 and **(d)** Step 2 in the nucleophile deprotonated, activated precursor (AP) state from MD simulations. Ions present in MD simulations are represented as spheres. 3D-RISM isodensity contours for the AP state of Step 1 where G_exo_ is **(e)** UCG and **(f)** guanosine (G). Mg^2+^ (magenta) binding densities are calculated from 3D-RISM and depicted at the 66^th^ contour level. Structures are depicted in parallel stereo with the surrounding ribozyme environment depicted as ribbons.

In contrast, site B exhibits step dependent behavior. Consistent with the cryo-EM densities, 3D-RISM predicts a stronger and more persistent population at site B in Step 2 (*ρ*_site_/*ρ*_bulk_ = 80) than in Step 1 (65), with density concentrated at the pro-*S_P_* oxygen of the scissile phosphate (Figure 2a,b). This difference suggests that the metal at site B, which coordinates a non-bridging oxygen, is more likely to be occupied in Step 2 than in Step 1. For reference, the density of site B in Step 1 is lower than for peripheral ions Mg(D) (76) and Mg(E) (68), but higher than Mg(D) (79) and Mg(E) (78) in Step 2. Isocontour values at positions near the active site are provided in Table S2, with detailed labels of the positions and coordinating phosphates shown in Table S1 and Figure S5 of the Supporting Information. An accompanying depiction of the 3D-RISM densities including the surrounding ribozyme environment is provided in Figure S4 of the Supporting Information.

To further interrogate this site, we considered the activated precursor (AP) in which the nucleophile is deprotonated. Upon activation, metal ion density generally increases across the active site, with the nucleophile-coordinating ion showing the highest occupancy (Figure 2c,d). The preferred location of site B shifts in Step 2 to a position between the pro-*S_P_* oxygen and the nucleophile, whereas in Step 1 it remains localized near the pro-*R_P_*oxygen. Note that the stereochemical pro-*R_P_*and pro-*S_P_*labels are reversed between steps, but they refer to the same NBO from a microscopic reversibility perspective. This redistribution indicates a potential change in binding mode upon nucleophile activation and highlights the need to distinguish between alternative coordination geometries.

Finally, the affinity of site B in Step 1 depends sensitively on the identity of the exogenous guanine cofactor. Substitution with UCG increases site B occupancy (77) relative to GTP (68), whereas guanosine decreases it (63), as shown in Figure 2e,f. These trends suggest that site B affinity is enhanced by the presence of a torsionally constrained, negatively charged phosphate at the G binding site, which creates an electrostatic environment resembling that of Step 2. Together, these results support a model in which site B is a conditionally populated metal binding site whose affinity and coordination mode are modulated by both the catalytic step and local electrostatics tuned by the exogenous cofactor.

### AFE simulations quantify active site Mg^2+^ binding affinities

Free energies were computed using MBAR, and *σ* is the uncertainty computed from the circular moving block bootstrap procedure described in ref 90. Binding free energies (ΔΔ*G_b_*) are relative to Mg^2+^ binding to a dinucleotide phosphate in solution (ref) (Figure S6), with negative values indicating more favorable binding in the ribozyme. AFE simulations were performed on the SS, with Mg(B) initially coordinating only the solvent-exposed non-bridging oxygen (NBO). GTP_exo_ is bound in Step 1.

We next quantified Mg^2+^ binding at sites within and peripheral to the active site (A–E) using alchemical absolute binding free energy (ABFE) simulations, in which each ion was independently annihilated across 16 *λ* windows by gradually decoupling its interactions with the surrounding environment. To isolate electrostatic contributions and prevent artifactual unfolding upon ion removal, we employed root mean square deviation (RMSD) restraints^93^ to maintain the active site geometry. Binding free energies (ΔΔ*G_b_*) are reported relative to the absolute binding affinity of Mg^2+^ coordinating a dinucleotide reference in solutions, and thus represent the affinity imparted by the THr environment relative to the reference. The resulting free energies (Tables 1 and S3) corroborate the 3D-RISM predictions, yielding a consistent affinity hierarchy of A *>* C *>* B across both catalytic steps. Formation of the activated precursor (AP) state generally strengthens binding at the coordinating site, reflecting enhanced electrostatic stabilization upon nucleophile deprotonation (Table S3).

Despite its weaker binding relative to sites A and C, site B exhibits affinity (−13.0 kcal·mol*^−^*^1^) comparable to the peripheral consensus sites Mg(D) and Mg(E) in Step 2 (−10.0 and −14.0 kcal·mol*^−^*^1^), indicating that it is likely to be populated for this step under experimentally relevant conditions. Notably, ABFE calculations show that site B is more favorable in Step 2 (−13.0 kcal·mol*^−^*^1^) than in Step 1 (−4.6 kcal·mol*^−^*^1^), in contrast to the stronger primary sites, highlighting its sensitivity to local electrostatic and structural context. Across simulations, Mg(B) predominantly coordinates the solvent exposed non-bridging oxygen, with dual coordination to both the non-bridging oxygen and nucleophile observed only in Step 2 in simulations where Mg(A) was removed.

Together, these results refine the solvation theory picture, establishing site B as a weaker but conditionally populated metal binding site, particularly in Step 2, whose affinity is modulated by catalytic state and local charge organization.

### MD simulations predict that Mg(B) enhances catalytic fitness in Step 2 (ligation)

Enzymes that catalyze phosphoryl transfer reactions employ four principal strategies — inline attack alignment, stabilization of negative charge on non-bridging oxygens (NBOs), activation of the nucleophile, and stabilization of the leaving group — termed *α*, *β*, *γ*, and *δ* catalysis, respectively.^39,105^ The interactions in THr that fulfill these criteria are summarized in Figure 3a. To define the role and preferred binding mode of Mg(B), we performed MD simulations of the standard state (SS; probing *γ* catalysis) and the activated precursor (AP; probing *α*, *β*, and *δ* catalysis) for Steps 1 and 2, with and without Mg(B) coordinated to the solvent facing NBO.

**Figure 3.**
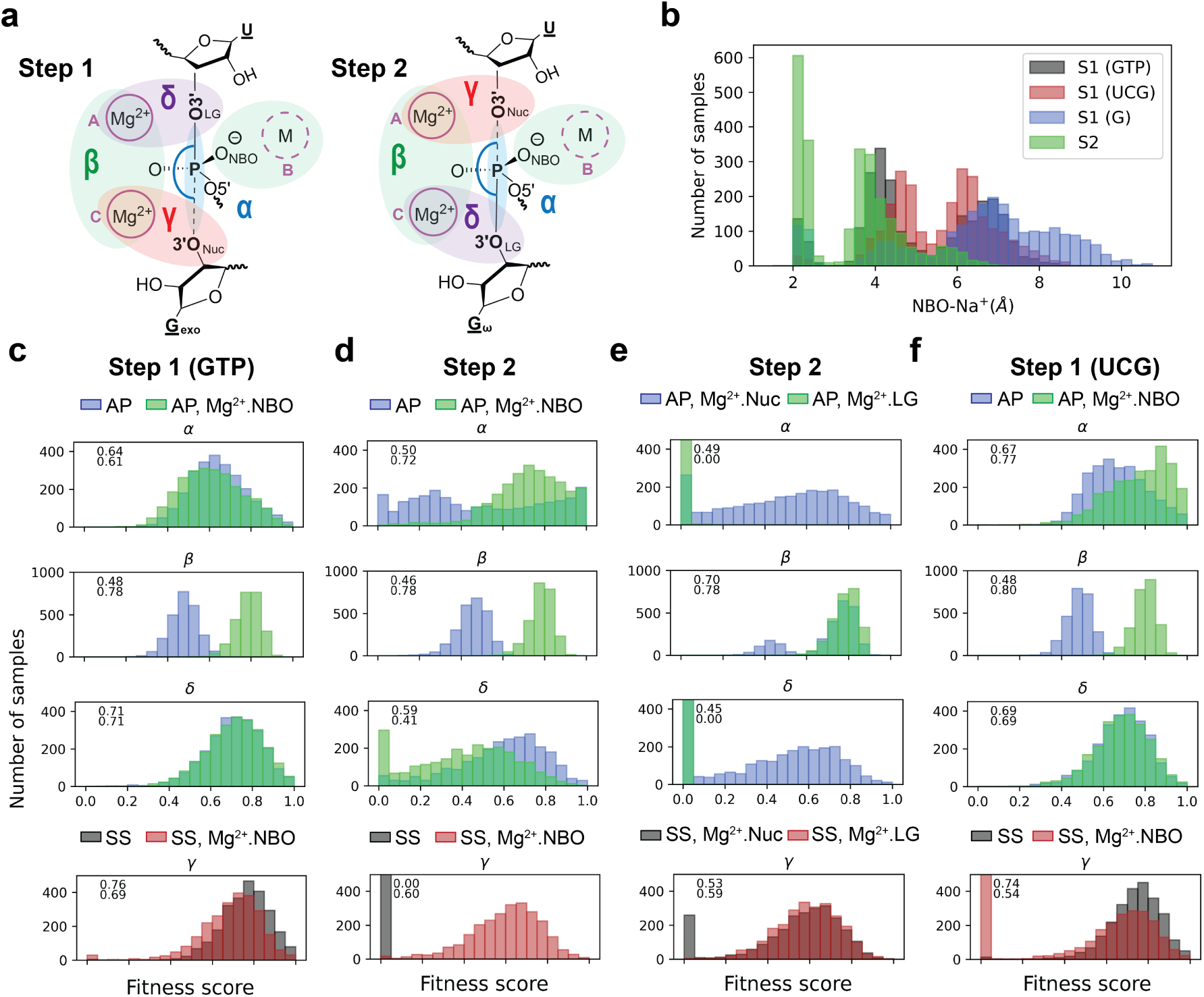
Impact of Mg(B) binding mode on inline fitness. **(a)** Roles of active site components that contribute to catalytic fitness in Step 1 (left) and Step 2 (right) of splicing. The inline attack distance is indicated by a dashed line, and angle by a blue arc. **(b)** Distribution of the closest Na^+^ ion distances to the solvent exposed NBO. Catalytic fitness of the standard state (SS, nucleophillic O3*^′^* protonated), and activated precursor (AP, nucleophilic O3*^′^* deprotonated) in the presence (Mg^2+^.NBO) and absence of Mg(B) for **(c)** Step 1 with GTP_exo_ and **(d)** Step 2. **(e)** Comparison of catalytic fitness of Step 2 with Mg(B) simultaneously coordinating the NBO and either the nucleophile (Nuc) or leaving group (LG). **(f)** Impact of Mg(B) on catalytic fitness in Step 1 with UCG_exo_.

In the absence of Mg(B), Na^+^ ions partially occupy site B and contribute to *β* catalysis, but their distribution is strongly step-dependent (Figure 3b). In Step 1, Na^+^ sampling around the NBO is diffuse and heterogeneous, whereas in Step 2 it predominantly occupies the first coordination shell, indicating a greater requirement for localized electrostatic stabilization. This behavior is consistent with the 3D-RISM and AFE results and reflects the increased negative charge density associated with the additional phosphate environment in Step 2.

Catalytic fitness distributions for all systems are shown in Figure 3c–f. Fitness criteria and system specific scores are provided in Tables S4–S5, and system specific fitness plots are are provided in Figures S7–S11 of the Supporting Information. Binding of Mg(B) universally enhances *β* catalysis, whereas *δ* catalysis is largely unaffected, consistent with a non-bridging coordination mode. By contrast, the impact on *α* and *γ* catalysis is step-dependent. In Step 1 (GTP-bound), *α* fitness is moderate in the absence of Mg(B) (0.64) and remains largely unchanged upon Mg(B) binding (0.61), while *γ* catalysis is high (0.76) and only modestly reduced (0.69). In Step 2, however, *α* fitness is initially moderate (0.50) but increases substantially upon Mg(B) binding (0.72). Moreover, *γ* catalysis is strongly suppressed in the absence of Mg(B) (0.0) due to the formation of a water mediated network that disrupts hydrogen bonding to O3*^′^*, but is restored upon Mg(B) binding (0.60) as this network is displaced. Altogether, these results indicate that Mg(B) is functionally more important in Step 2 than in Step 1 for enhancing *α*, *β*, and *γ* catalysis.

To resolve the binding mode of Mg(B), we tested alternative geometries in which the ion bridges the NBO with either the nucleophile or the leaving group in Step 2 (Figure 3e). In these configurations, Mg(B) consistently disrupts inline (*α*) fitness, thereby precluding them as functionally important. While coordination with the nucleophile yields modest *α* fitness (0.49), the binding mode in 1 replica reverts back to indirect coordination of O3*^′^*, and *α* fitness is higher (0.72) for this mode. In contrast, simulations in which Mg(B) is initiated indirectly coordinated to O3*^′^* never migrate to direct coordination. Overall, the data support a model in which Mg(B) directly coordinates only the solvent-exposed NBO.

Finally, the dependence of catalytic fitness on the exogenous guanine cofactor in Step 1 (Figure. 3f, Table S5) further clarifies the role of Mg(B). Moderate *α* fitness is maintained with G, GTP, or UCG in the absence of Mg(B), indicating that cleavage is intrinsically robust. However, UCG exhibits enhanced fitness upon Mg(B) binding (0.77), consistent with the formation of a more defined, Step 2-like electrostatic environment due to its additional, torsionally constrained phosphate.

In all simulations, we observe consistent coordination between the O2*^′^* of G_exo_ or G*_ω_* with Mg(C) consistent with experiment.^76^ Substitution of the O2*^′^* by OCH_3_ has been shown to reduce activity, whereas NH_2_, and NHCH_3_ retained activity in the presence of Mn^2+^.^106^ This suggested the HO2*^′^* of G donates a hydrogen bond that is critical for cofactor binding,^106^ but the identity of the hydrogen bond acceptor was unclear. As shown in Figure S12, the HO2*^′^* predominantly hydrogen bonds to the phosphate of residue 307, with transient interaction with the phosphate of residue 262 that interrupts the Mg(C) coordination sphere.

Together, MD simulations indicate that Mg(B) is not required for catalytic fitness in Step 1, but becomes functionally important in Step 2, where it stabilizes the increasingly strained electrostatic environment while maintaining a binding mode restricted to the coordination of the solvent facing NBO.

### QM/MM+ΔMLP free energy simulations suggests different metal ion requirements between Steps 1 and 2

Having established an active state ensemble and plausible binding modes for Mg(B), we next evaluated the reaction mechanism using QM/MM+ΔMLP free energy simulations. Experimental activity-pH profiles of THr indicate an increasing log-linear dependence with pH with an apparent p*K*_a_ greater than 13,^108–110^ consistent with nucleophile deprotonation as the dominant ionization event. We therefore examined whether additional general acid/base contributions are required, and whether Mg(B) or Na(B) could participate via a coordinated water.

Free energy profiles were computed using umbrella sampling with a QM/MM Δ-machine learning potential (ΔMLP). Specifically, we use a range-corrected equivariant graph neural network based on the MACE architecture,^111^ trained to reproduce PBE0/6-31G* energetics from a DFTB3/3ob reference (QM/MM+ΔMLP).^89,112,113^ This approach enables efficient sampling of *ab initio* quality free energy surfaces. Additional details regarding training data generation and validation of the model on training and test sets are given in Figures S13-S15 and Table S6 of the Supporting Information.

#### Step 1

In the absence of Mg(B), multiple protonation pathways for leaving group departure were considered as indicated by blue arrows in Figure 4a. All general acid mechanisms, including proton transfer from water or O2*^′^*, are disfavored, yielding barriers (*>*24 kcal·mol*^−^*^1^) well above that of direct oxyanion departure (21.6 kcal·mol*^−^*^1^). Free energy profiles and associated barriers for energetically unfavorable mechanisms can be found in Figure S16 and Table S7 of the Supporting Information. These results, together with experimental substituent effects,^114^ indicate that Step 1 likely proceeds without general acid catalysis, with Mg(A) functioning primarily as a Lewis acid.

**Figure 4.**
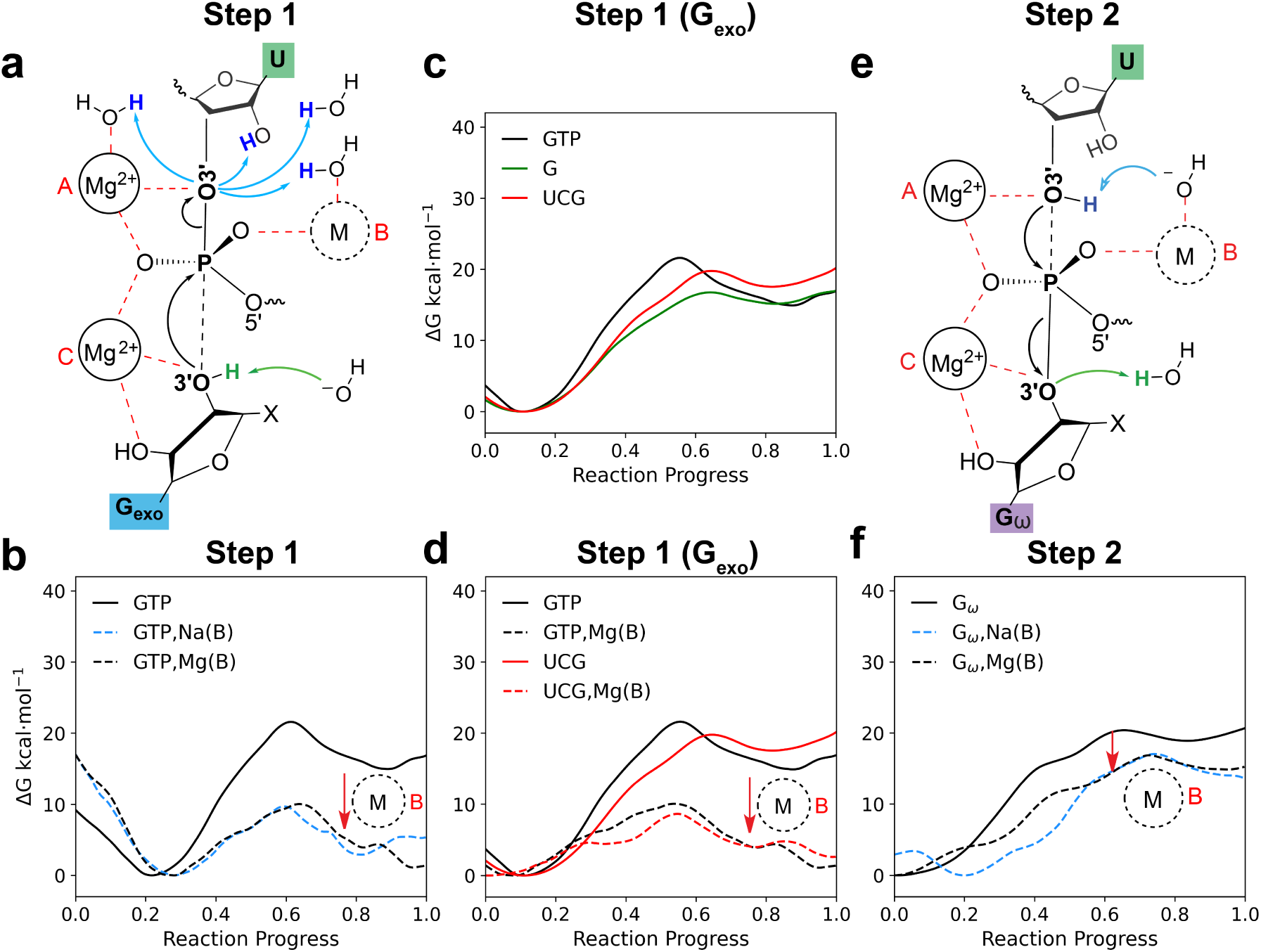
Free energy profiles from ΔMLP accelerated QM/MM simulations of Step 1 and 2 of splicing **(a)** Mechanistic scenarios probed by QM/MM+ΔMLP simulations in Step 1. Mg^2+^ ions A and C (marked in red) are always present, while M(B) (dotted circle) is either Mg^2+^, Na^+^, or not present. Phosphoryl transfer is denoted in black, nucleophile deprotonation in green, and leaving group protonation in blue. Metal ion coordination is indicated by red dashed lines. **(b)** Free energy profiles of Step 1 with GTP_exo_. Free energy profiles of Step 1 with G_exo_ being either G, GTP, or UCG in the **(c)** absence or **(d)** presence of Mg(B). **(e)** Mechanistic scenarios probed in Step 2. **(f)** Free energy profiles of Step 2. The G*_ω_* system contains a QM Na^+^ indirectly coordinating the NBO. The general reduction in free energy barrier induced by M(B) directly coordinating the NBO is indicated by a red arrow.

In addition, deprotonation of the nucleophile by a hydroxide indirectly coordinating Mg(C) (the only clear hydrogen bonding partner of HO3*^′^*) is exoergic, supporting pre-reaction activation of the nucleophile by a specific base in accord with the log-linear dependence of the activity-pH profile in the range of 5.5-9. However, even the most favorable barrier in the absence of Mg(B) (21.6 kcal·mol*^−^*^1^ in Figure 4b and Table 2) remains above the experimental estimate of 16–17 kcal·mol*^−^*^1^.^115,116^

**Table 1.** Relative binding free energies of active site Mg^2+^ ions in THr from AFE simulations.

| | $\Delta\Delta G_b = \Delta G_b(\text{THr}) - \Delta G_b(\text{ref})$ (kcal·mol<sup>-1</sup>) | | | | |
| --- | --- | --- | --- | --- | --- |
| Step | A | B | C | D | E |
| 1 | -41.6±0.6 | -4.6±0.7 | -18.8±0.4 | -17.3±1.3 | -18.1±1.4 |
| 2 | -49.4±2.2 | -13.0±1.1 | -25.5±3.2 | -10.0±0.9 | -14.0±0.9 |

**Table 2.** Free energy barriers of phosphoryl transfer reactions and associated properties for mechanisms studied with QM/MM+ΔMLP simulations.

|  |  |  | kcal·mol<sup>-1</sup> |  | Å | kcal·mol<sup>-1</sup> |  | min<sup>-1</sup> |
| --- | --- | --- | --- | --- | --- | --- | --- | --- |
| System | Acid | Base | $\Delta G^\ddagger$ | $\sigma$ | $\xi_{PT}$ | $\Delta G_{rxn}$ | $\sigma$ | $k_{int}$ |
| Step 1 |  |  |  |  |  |  |  |  |
| GTP | - | OH<sup>-</sup> | 21.59 | 0.17 | 0.99 | 14.98 | 0.31 | 5.47x10<sup>-2</sup> |
| GTP,Na(B) | Na(B) | OH<sup>-</sup> | 9.72 | 0.10 | 0.43 | 5.28 | 0.17 | 2.77x10<sup>7</sup> |
| GTP,Mg(B) | Mg(B) | OH<sup>-</sup> | 10.05 | 0.07 | 0.33 | 1.10 | 0.12 | 1.59x10<sup>7</sup> |
| G | - | - | 16.76 | 0.08 | 0.46 | 15.13 | 0.12 | 1.90x10<sup>2</sup> |
| UCG | - | - | 19.78 | 0.09 | 0.55 | 17.54 | 0.08 | 1.16 |
| UCG,Mg(B) | Mg(B) | - | 8.64 | 0.09 | 0.65 | 2.59 | 0.42 | 1.72x10<sup>8</sup> |
| Step 2 |  |  |  |  |  |  |  |  |
| G<sub>ω</sub> | - | -* | 20.38 | 0.33 | 0.63 | 18.90 | 0.30 | 4.22x10<sup>1</sup> |
| G<sub>ω</sub>,Na(B) | - | Na(B) | 17.05 | 0.09 | 0.47 | 13.64 | 0.15 | 1.17x10<sup>2</sup> |
| G<sub>ω</sub>,Mg(B) | - | Mg(B) | 16.87 | 0.12 | 0.70 | 14.88 | 0.18 | 1.58x10<sup>2</sup> |

$\Delta G^\ddagger$ indicates the rate controlling free energy barrier. Standard error estimates ( $\sigma$ ) were computed from bootstrapping as described in ref 107. The reaction coordinate ( $\xi_{PT}$ ) corresponds to the phosphoryl transfer coordinate at the transition state. Positive values indicate the leaving group phosphorous distance exceeds the nucleophile phosphorus distance. Intrinsic reaction rates ( $k_{int}$ ) were estimated from Equation 6 using $\Delta G^\ddagger$ . A (-) symbol indicates the absence of acid or base. A (\*) indicates a system where Na<sup>+</sup> indirectly coordinates the solvent facing NBO at the B site, but does not play an acid or base role. Metals in an acid or base role act through either a water or hydroxide, respectively.

Inclusion of a cation at site B substantially lowers the barrier. Both Na^+^ and Mg^2+^ reduce the barrier to ∼10 kcal mol*^−^*^1^ (Figure 4b), largely by stabilizing negative charge at the NBO and facilitating leaving group departure (*β* and *δ* catalysis, respectively). In addition, nucleophile deprotonation becomes more favorable, indicating a lowering of the O3*^′^* p*K*_a_. Notably, when Mg(B) is present, proton transfer from a coordinated water becomes favorable, indicating a viable general acid pathway under these conditions. However, MD and 3D-RISM results indicate that site B is only weakly and transiently occupied in Step 1 with GTP_exo_, implying that this low barrier pathway is not always accessible. Thus, Step 1 likely proceeds predominantly via Lewis acid catalysis, with transient occupation of site B by either Na^+^ or Mg^2+^ providing additional stabilization and p*K*_a_ tuning under favorable conditions when GTP is bound.

#### Cofactor and electrostatic effects

The identity of the exogenous guanine cofactor further modulates these trends. In Step 1, barriers with GTP or UCG exceed those with G in the absence of Mg(B), reflecting electrostatic repulsion from additional phosphate groups (Figure 4c,d). While G exhibits an experimentally consistent barrier without M(B) (16.8 kcal·mol*^−^*^1^), binding of Mg(B) alleviates the electrostatic penalty for GTP and UCG, yielding barriers below those of G (10.1 and 9.0 kcal·mol*^−^*^1^, respectively). This suggests enhanced Mg(B) affinity for cofactors with torsionally constrained phosphates, consistent with 3D-RISM and MD results. Experimentally, G analogs including UCG demonstrate faster binding kinetics than G due to base pairing with the P9 duplex, but the intrinsic reaction barriers are expected to be similar,^110,117^ perhaps achieved through differential transient interactions with Mg(B). Finally, the reverse reaction of Step 1 has often been used as an analog for probing Step 2,^109^ but the differences in barriers resulting from different Step 1 electrostatic environments shown here also highlight that this assumption neglects potential differences between G_exo_ and G*_ω_*.

#### Step 2

The barrier for Step 2 is also sensitive to occupation of site B (Figure 4e,f). When Na^+^ or Mg^2+^ directly coordinates the NBO, the barrier (∼17 kcal·mol*^−^*^1^) closely matches experimental estimates. In this binding mode, Mg(B) can also act as a base via a coordinated hydroxide to deprotonate O3*^′^*, as shown in Figure 4e. Outersphere coordination of Na^+^ to the NBO increases the barrier by *>*3 kcal·mol*^−^*^1^, indicating that effective catalysis requires direct, persistent binding at site B. Combined with catalytic fitness, 3D-RISM, and ABFE results, this suggests that site B is consistently occupied in Step 2, in contrast to its transient occupation in Step 1. As shown in Figure 5a,c the transition state structures of Step 1 and 2 experience inherently different leaving group stabilization, with HO2*^′^* of U(−1) forming an additional stabilizing hydrogen bond to the leaving O3*^′^* in Step 1, but not Step 2. This is reflected in higher product energy in Step 2 (Figure 5b,d). Another fundamental difference between Step 1 and 2 arises from the fact that Mg(A) and Mg(C) coordinate net charges of −4 and −3 respectively from phosphates, with Mg(C) and Mg(D) bridged by one of these phosphates. Nucleophile formation in Step 2 increases the charge around Mg(A) to −5, likely increasing the need for activation by Mg(B) in Step 2 relative to Step 1.

**Figure 5.**
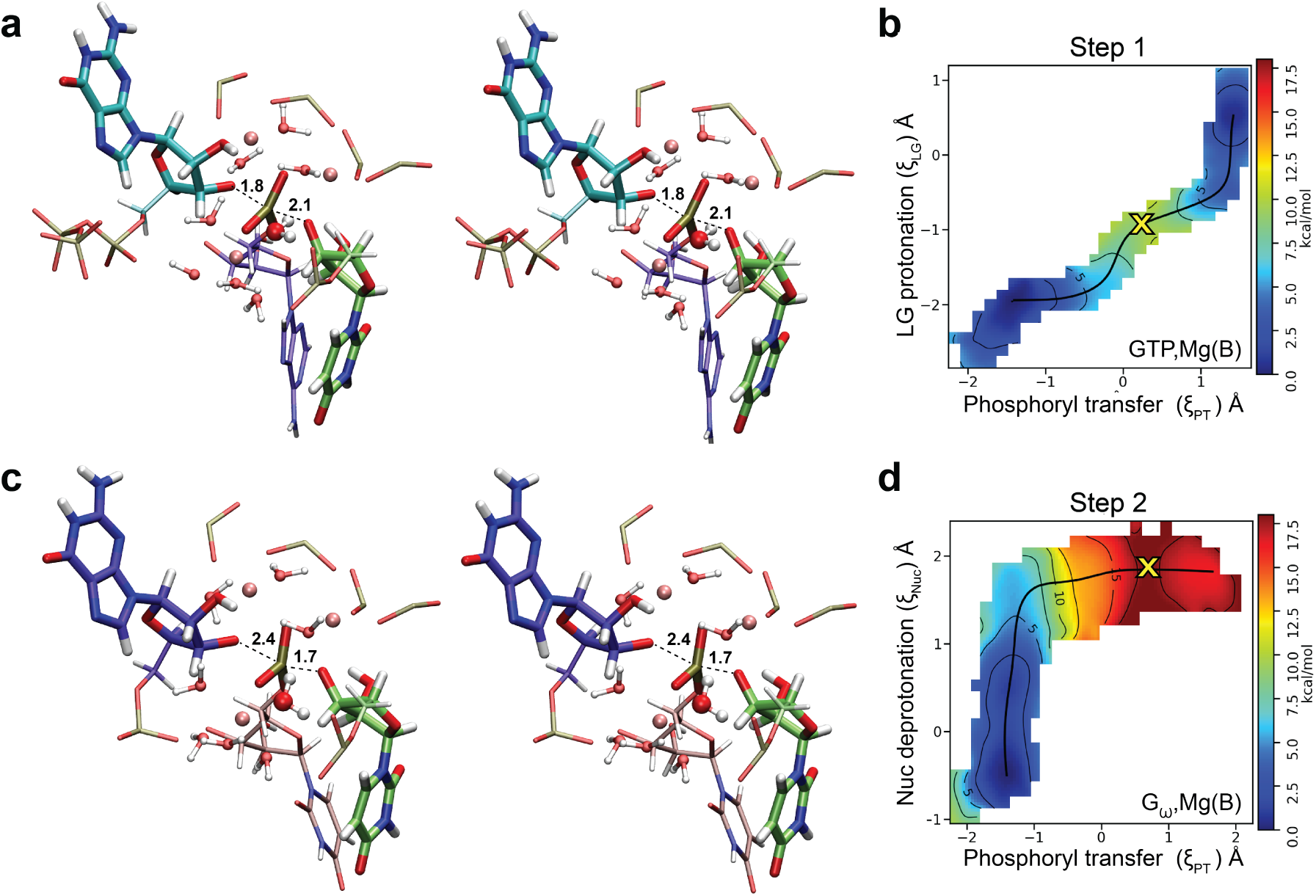
Transition state structures and free energy surfaces from ΔMLP accelerated QM/MM simulations. **(a)** Representative transition state structure in the presence of Mg(B) for Step 1 with GTP_exo_. Structures are depicted in parallel stereo with mean transition state phosphoryl transfer distances provided in Å. **(b)** Corresponding 2 dimensional free energy surface for Step 1. Reaction coordinate definitions are given in Equation 3, and the transition state is marked with an “x”. **(c)** Representative transition state structure in the presence of Mg(B) for Step 2. **(d)** Corresponding 2 dimensional free energy surface for Step 2.

In summary, the free energy profiles and complementary simulations support a unified model in which Step 1 proceeds primarily via Lewis acid catalysis by Mg(A), with only transient or fractional occupation of site B by Na^+^ or Mg^2+^ when G_exo_ carries a phosphate. In contrast, Step 2 requires near complete occupation of site B to achieve experimentally consistent barriers, which MD simulations suggest would not be achieved with Na^+^ alone. This divergence is consistent with experimental Mg^2+^-dependence, in which the rate of Step 1 increases continuously with Mg^2+^ concentration,^118^ whereas Step 2 saturates at low millimolar concentrations.^119^ Together, these results establish that the two steps of splicing have fundamentally different metal ion requirements, with divalent or monovalent ions playing a cofactor dependent, transient role in Step 1, but a persistent and functionally essential role in Step 2.

### Isofunctional substitution of a conserved G·U wobble maintains key hydrogen bonding network to expand targets for splicing

The ultimate goal of computational enzymology is often either to guide the inhibition of enzymes or to enhance catalysis when enzymes are deployed as biotechnological tools. THr is relevant to both contexts. Group I introns occur in fungi, bacteriophages, protozoa, and eubacteria,^1^ making them potential targets for inhibition by small molecules.^120,121^ Structure based screening has already enabled the discovery of inhibitors targeting related introns, demonstrating the therapeutic potential of RNA directed small molecules.^122–124^ Accurate modeling of metal ion interactions is essential in this context because divalent ions strongly influence RNA structure and ligand binding energetics.^125^

At the same time, THr has emerged as a powerful platform for RNA editing through trans-splicing, with potential therapeutic applications including sickle-cell disease,^126^ myotonic dystrophy,^127^ and several cancers.^3,128–131^ In these approaches, sequence specificity can be programmed by tuning residues of the internal guide sequence (IGS) to base pair with a target RNA. A key limitation, however, is the requirement that the nucleotide immediately upstream of the splice site [position (–1)] be U, which forms a conserved G22·U wobble pair to the IGS within the active site (Fig. 6a). In the first step of splicing, this U(–1) becomes the leaving group and later the nucleophile in the second step of splicing. Experiments have shown that a G·C(–1) pair can be tolerated only at low pH, where protonation of cytosine forms a G·C^+^ wobble pair; at neutral pH the interaction shifts toward Watson–Crick geometry and substrate docking is reduced by approximately two orders of magnitude (Fig. 6b,c).^132^ The interactions made by the exocyclic amine of G22 are expected to contribute at least 2 kcal·mol*^−^*^1^ to substrate binding,^133^ and an additional 1 kcal·mol*^−^*^1^ to the transition state relative to an inosine·U pair.^134^ 2,6-diaminopurine at position 22 (diAP) has been shown to be highly active with C(−1), but the rate of miscleavage is about 20 fold higher than G·U.^133,134^ If cytosine could instead be accommodated at near neutral physiological pH (7.0-7.4) with high fidelity, the range of potential RNA targets for therapeutic trans-splicing would expand substantially.

**Figure 6.**
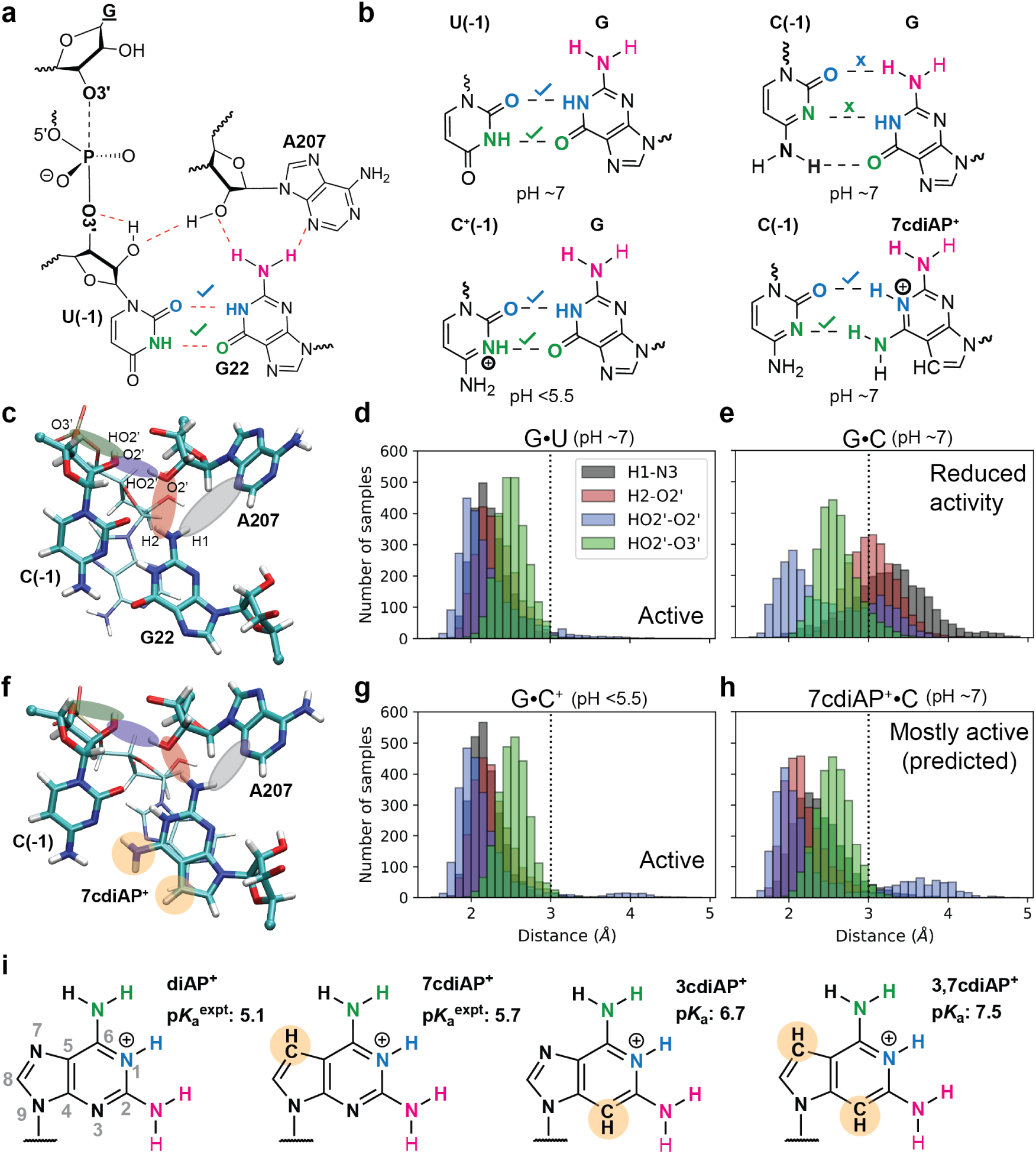
Mutation of the conserved active site G·U wobble base pair upstream of the splice site in Step 1. **(a)** Key hydrogen bonding network involving the conserved active site G·U. **(b)** Base pairing interactions formed by the simulated nucleotides. **(c)** Representative MD simulated structures of the G·U replaced by G·C. Histograms quantifying the hydrogen bond distances in the **(d)** G·U and **(e)** G·C. **(f)** Representative MD simulated structures of the G·U replaced by 7cdiAP^+^·C. Atomic substitutions in 7cdiAP^+^ relative to G are indicated in tan. Histograms quantifying the hydrogen bond distances in the **(g)** G·C^+^ and **(h)** 7cdiAP^+^·C containing systems. Colors match those of the hydrogen bonds highlighted in **c** and **f**. A black vertical line at 3 Å indicates the tail distance for a hydrogen bond. **(i)** Structures of modified diAP nucleotides with experimental or predicted p*K*_a_s proposed to be capable of pairing with C(−1).

Structural and computational analyses indicate that the functional requirement for a wobble pair arises from its role in organizing a hydrogen bonding network that stabilizes the docked substrate. In the native complex, the exocyclic amine of G22 hydrogen bonds to N3 and O2*^′^* of A207, and the O2*^′^* of A207, in turn, hydrogen bonds to the O2*^′^* of the (–1) nucleotide (Figure 6a). This relay directs the formation of a stabilizing hydrogen bond to the leaving group oxygen and helps organize the active site for catalysis.^46,70,132,135,136^ When cytosine replaces uracil at neutral pH, the G·C Watson–Crick geometry disrupts this network (Figure 6c), resulting in the observed loss of docking efficiency. At low pH (*<* 5.5), however, protonation of cytosine restores a wobble-like G·C^+^ interaction that preserves these contacts and supports catalysis (Figure 6b).^132^

These observations led us to hypothesize that an isofunctional substitution at position 22 might allow cytosine to be accommodated at neutral pH while preserving the essential hydrogen bond network. We therefore sought to design a substitution of G22 with atomic variants of 2,6-diaminopurine (diAP, p*K*_a_ 5.1^137^) that would retain the exocyclic amines and tune the p*K*_a_ at the N1 position to enable a wobble-like pair to form with C(−1) as shown in Figure 6b. It should be noted that neither the N7 or N3 positions of G22 are observed to be involved in important structural hydrogen bonds with the ribozyme or substrate in any of the available experimental structures or the simulations presented here. We therefore first consider 7-deazapurine-2,6-diamine (7cdiAP), which has been synthesized and tested, and exhibits an elevated N1 p*K*_a_ of 5.7 in solution.^138–140^ As the apparent p*K*_a_ of C(–1) in the G22·C(−1)^+^ pair in the ribozyme is shifted ∼1.5 units from 4.5 to ∼6,^132^ we anticipate a similar p*K*_a_ shift at the N1 position of 7cdiAP when in a wobble-like pair with C(−1). This would imply that the N1 protonated form, 7cdiAP^+^, could remain populated near neutral pH and thus stabilize a functional wobble-like interaction, as illustrated in Figure 6b,f.

Molecular dynamics simulations support this hypothesis (Table 3). Replacing the native G·U pair with a neutral G·C pair shifts the base pairing toward Watson–Crick geometry and disrupts hydrogen bonds to A207, destabilizing the O2*^′^*-mediated network (Figure 6c,d,e). In contrast, simulations of G·C^+^ maintain these interactions, consistent with the experimentally observed tolerance at low pH (Figure 6g). Substitution with 7cdiAP^+^ paired to C preserves the key hydrogen bonds and restores the geometry of the interaction network relative to the G·C pair (Figure 6f,h). These trends persist in simulations performed both in the absence and presence of Mg(B) (Table 3 and Figure S17 of the Supporting Information), indicating that the modified base maintains the contacts that anchor the substrate during docking.

**Table 3.** Mean and standard deviation of key hydrogen bond distances associated with wobble base pair variants.

| Nucleotide |  | Distance (Å) |  |  |  |
| --- | --- | --- | --- | --- | --- |
| 22 | -1 | H1-N3 | H2-O2' | HO2'-O2' | HO2'-O3' |
| Mg(B) not present |  |  |  |  |  |
| G | U | 2.24 (0.23) | 2.33 (0.25) | 2.21 (0.38) | 2.52 (0.19) |
| G | C | 3.27 (0.50) | 3.04 (0.31) | 2.48 (0.55) | 2.61 (0.25) |
| G | C<sup>+</sup> | 2.19 (0.19) | 2.27 (0.25) | 2.18 (0.44) | 2.53 (0.19) |
| 7cdiAP<sup>+</sup> | C | 2.41 (0.35) | 2.21 (0.28) | 2.41 (0.72) | 2.57 (0.22) |
| Mg(B).pro-$R_P$ | | | | | |
| G | U | 2.22 (0.21) | 2.31 (0.24) | 2.18 (0.39) | 2.56 (0.18) |
| G | C | 3.45 (0.54) | 3.22 (0.34) | 2.38 (0.54) | 2.61 (0.25) |
| G | C<sup>+</sup> | 2.18 (0.20) | 2.22 (0.21) | 2.16 (0.24) | 2.60 (0.18) |
| 7cdiAP<sup>+</sup> | C | 2.34 (0.38) | 2.40 (0.49) | 2.15 (0.32) | 2.49 (0.21) |

Ideally, an isofunctional diAP mutant at position 22 would have complete occupancy of the N1 proton at neutral pH to pair with C(−1). Whereas the 7-deaza modification in 7cdiAP up-shifts the p*K*_a_ of diAP by 0.6 units, deaza modification at the N3 position is expected to up-shift the p*K*_a_ even further. Experimentally, adenosine has a p*K*_a_ of 3.6 at the N1 position, whereas 7-deazaadenosine and 3-deazaadenosine have p*K*_a_ values of 5.3 and 6.8, respectively.^141^ Given that the N3 position of G22, like the N7 position, does not form any functionally crucial hydrogen bonds, this position could also be a candidate for modification, although we were not able to find literature outlining the synthesis and testing of 3-deazapurine-2,6-diamine (3cdiAP) as we did for 7cdiAP. Nonetheless, we performed high-level quantum chemistry calculations with implicit solvation effects to predict the p*K*_a_ values for chemically modified diAP variants 3cdiAP (∼6.7) and 3,7cdiAP (∼7.5) (see Figures S18-S19 and Table S8-S10 of Supporting Information for details). Based on these results, we predict that 3cdiAP and 3,7cdiAP will have even higher occupancy of the functional protonated forms in the physiologic pH range (Figure 6i).

Together, these results highlight the structural role of the conserved wobble pair in organizing the catalytic hydrogen-bonding network of THr and suggest that targeted nucleobase substitutions can be used to modulate this interaction. In particular, substitution of G22 with 7cdiAP, 3cdiAP and 3,7cdiAP appear capable of maintaining the active site architecture while accommodating C(–1) at neutral pH, providing a potential strategy for expanding the sequence space accessible to programmable trans-splicing ribozymes.

## Conclusion

In this work we have used a combination of 3D-RISM, ABFE, MD, and ΔMLP accelerated QM/MM simulations to investigate catalytic requirements of THr in the first and second step of trans-splicing. Our simulations suggest a potential interplay between a monovalent or divalent metal ion binding at the solvent exposed NBO, with affinity dependent on the identity of G_exo_. We rule out the possibility of a third Mg^2+^ ion binding the nucleophile, as well as mechanistic pathways involving water or O2*^′^* acting as an acid, as well as involvement of a specific base. When the nucleophile is G in Step 1, an oxyanionic leaving group yields an experimentally consistent barrier of 16.76 kcal·mol*^−^*^1^, but when GTP or UCG are present, Na^+^ or Mg^2+^ are required at the B site to access an experimentally feasible barrier, likely with fractional occupancy. In Step 2, Na^+^ or Mg^2+^ are required with high occupancy to achieve experimentally consistent barriers of 17.05 kcal·mol*^−^*^1^ and 16.87 kcal·mol*^−^*^1^, respectively. Finally, we propose and computationally evaluate modifications to G22 — specifically, isofunctional substitutions of 2,6-diaminopurine with tuned p*K*_a_ values (7cdiAP, 3cdiAP and 3,7cdiAP) — that are predicted to enable accommodation of C(–1) in a critical wobble-like base pair at physiological pH. This design offers a potential route to expand the sequence space accessible to programmable trans-splicing ribozymes for RNA editing applications. Altogether, our findings have implications for both structure based drug design efforts and the design of therapeutic trans-splicing ribozymes. Our findings also highlight an intriguing asymmetry between the first and second steps of the reaction, potentially suggesting an important design principle for ribozymes capable of splicing.

## Materials and Methods

In the present work, we apply an integrated computational enzymology approach to investigate the cleavage and ligation mechanisms of THr at *ab initio* DFT level accuracy. Specifically, our approach combines statistical mechanical solvation theory (3D-RISM) and alchemical free energy (AFE) simulations to identify functional divalent metal ion binding modes, long-time classical molecular dynamics (MD) simulations to assess catalytic fitness and characterize the catalytically active state in solution, and quantum mechanical/molecular mechanical + machine learning potential (QM/MM+ΔMLP) simulations to identify minimum free energy pathways corresponding to different mechanistic scenarios. Classical molecular dynamics and alchemical free energy simulations were performed with the GPU-accelerated pmemd.cuda engine within AMBER,^93,142^ 3D-RISM calculations were performed with RISM3D^99,100^ in AmberTools,^143^ and QM/MM+ΔMLP simulations were conducted using interoperable software infrastructure^144,145^ implemented in sander.MPI with advanced free energy analysis and path methods in FE-ToolKit.^90^

### System preparation

Three main system builds were prepared for the L-16 S*ca*I *Tetrahymena* ribozyme departing from the recent cryo-EM structures.^45,46^ Build 1, the pre-catalytic state of the cleavage step (Step 1),^46^ was built from PDB ID 7YCH.^46^ Build 2, like 1, was built from 7YCH with the addition of a UC before the G_exo_ (UCG_exo_), where the leading UC was modeled by aligning the nucleobase of G*_ω_* from Step 2 onto the nucleobase of GTP_exo_. Build 3, the exon ligation step (Step 2)^45^ was built from the cryo-EM structure 7YGA.^45^ In all, sulfur substitutions at the scissile phosphates were replaced by oxygen.

Cryo-EM structures of the first and second step contained 3 and 8 resolved Mg^2+^ ions, respectively. Additional Mg^2+^ ions were aligned onto both structures from PDB ID 7EZ2,^44^ resulting in each main build having 31 Mg^2+^. The three builds were each solvated with a truncated octahedron TIP4P-Ew water box,^146^ neutralizing Na^+^, and additional NaCl to yield a physiological concentration of 140 mM, yielding 70k waters and 300k particles (∼140 Å lattice vectors). The rest of the systems tested were obtained by making slight modifications to these three main builds. Box contents of each build and their variants are listed in Table S11 of the Supporting Information.

All systems were relaxed in two stages. The first stage focuses on equilibrating the solvent during which the solute molecules are tightly restrained, and the second stage slowly relaxes the molecular restraints in a manner of halving the force constants at each step. Specifically, the first stage starts with density equilibration, which involves a 500-step minimization, followed by 5 ps + 5 ps + 190 ps NPT simulations. These short simulations help accommodate large changes in the box size when density is equilibrating. Then another 500-step minimization resets velocities, and NVT heating from 0 K to 300 K over 600 ps slowly brings the system up to speed which is then held at 300 K for 1 ns. The first stage concludes with a 5 ns NPT simulation to allow for solvent relaxation. Throughout the first stage the molecule is restrained with a force constant of 50 kcal· mol^-1^·^Å-2^. The second stage starts with reducing it to 25 kcal· mol^-1^·^Å-2^. To accommodate for this large change in forces another 500-step minimization, and 1.6 ns NVT heating steps are carried out. Then 4 sets of 500 ps NPT simulations with force constants 25, 10, 5, and 2 kcal· mol^-1^·^Å-2^, respectively, finalize the relaxation of the molecule in solution.

Variants were generated by making small adjustments to the corresponding main build, then equilibrated with short minimization and heating steps to allow the solvent environment to adjust to the changes. Any net charge introduced by the modifications was neutralized by removing Na^+^ or Cl*^−^* ions as needed. Mg(B) was introduced by converting a Na^+^ to Mg^2+^. The guanosine containing system was built by replacing the phosphate tail of GTP with OH.

### Molecular mechanical force fields

The ribozyme was described with the ff99OL3 force field.^147,148^ The solvent environment was described with TIP4P-Ew water model,^146^ with Joung and Cheatham monovalent ion parameters^149^ and the Li and Merz 12-6-4 divalent ion parameters^150^ with Panteva corrections^151^ which balance interactions between Mg^2+^ and nucleic acids. Charges for the non-standard residues (GTP, GTP*^−^*, G*^−^*, U*^−^*, C^+^, 7cdiAP^+^) were determined using the restrained electrostatic potential (RESP) charge fitting method,^152,153^ with potentials calculated at HF/6-31G* level.

### 3D-RISM and Poisson–Boltzmann calculations

Single point three-dimensional reference interaction site model (3D-RISM)^99,100^ calculations were performed on cryo-EM structures and MD-derived conformations to identify candidate Mg^2+^ binding sites that may influence the catalytic mechanisms of THr. Systems were stripped of solvent and ions including Mg^2+^. DRISM was performed as described in ref 38 with the rism1d program to compute site-site solvent susceptibilities for a solution of 10 mM Mg^2+^, 140 mM Na^+^, and 160 mM Cl*^−^* (32,768 grid points, 0.025 Å spacing). The modified direct inversion of the iterative subspace (MDIIS) approach was used to iteratively solve the DRISM equation with PSE2 closure to a residual tolerance of 10*^−^*^12^ at 298 K and a dielectric constant of 78.497 for bulk water. A constant density approach was used in which the density of water was assumed to be 55.296 M using the SPC/E water model.^154^ Single point 3D-RISM was performed on a 320×320×320 ^Å3^ grid with a 160×160×160 ^Å3^ solvation box without solvent-solvent interaction cutoff. PSE1 closure was employed to a residual tolerance of 10*^−^*^4^. A Poisson–Boltzmann (PB) calculation was performed on the pre-cleavage structure (7YCH). The pbsa solver^94–96^ was used to output the spatial distribution of the electrostatic potential. The grid spacing for the finite difference PB solver was set at the default value of 0.5 Å, and the dielectric interface was implemented with a level set function and the revised density function approach (ipb = 2, sasopt = 2).

### Alchemical free energy simulations

Alchemical free energy simulations were used to estimate binding affinity trends of active site Mg^2+^ ions at sites A, B, C, D, and E. An absolute binding free energy (ABFE) approach was taken by annihilating each Mg^2+^ ion independently in the contexts of the standard state (nucleophile protonated) and activated precursor (nucleophile deprotonated) with Mg(B) present, totaling 20 states. Root mean square deviation (RMSD) restraints^93^ relative to the average structure from production MD (100 ns) for each state were applied to the active site phosphates (C208, C262, A304, A306) with restraining force constants of 5 kcal·mol*^−^*^1Å^*^−^*^1^. Boresch style restraints^155^ were applied in a *λ* dependent fashion to the Mg^2+^ atom being eliminated to maintain the dummy atom’s position in the binding pocket. These included the distance from the Mg^2+^ ion to the pro-*S_P_* of the scissile phosphate (Mg^2+^-pro-*S_P_*), the angle between the Mg^2+^ ion, pro-*S_P_*, and phosphorous (Mg^2+^-pro-*S_P_* -P), and the torsion between the Mg^2+^ ion, the pro-*S_P_*, phosphorous, and pro-*R_P_* (Mg^2+^-pro-*S_P_* -P-pro-*R_P_*), with stereochemistry described according to the cleavage reaction. The values of the restraints and associated force constants were computed from production MD simulations. The recently developed smooth-step softcore potentials^156^ were used to create intermediate states that connect the real and dummy states. For each trial, the equilibration process included 1 ns of free MD on the end state followed by sequential 5000-step energy minimizations to generate initial *λ* window structures. Each *λ* window was then heated from 200 to 298 K over 100 ps to initiate velocities, and equilibrated in the NTP ensemble for an additional 100 ps. Production sampling was performed for 5 ns in the NTP ensemble with a 4 fs time-step. Hamiltonian replica exchange^157–160^ attempts were made, and BAR statistics were written, every 400 fs. The *λ* schedule was initially optimized from a so called “burn in” simulation of 32 linearly spaced *λ* windows for 0.8 ns for the annihilation of Mg(A) in the standard state of Step 1. The *λ* schedule was obtained by performing a 2 parameter optimization of the soft-core scaling (SSC) function to optimize the replica exchange acceptance ratio with 16 *λ* windows,^92^ yielding the following schedule: 0.00000, 0.13854, 0.21369, 0.27510, 0.32981, 0.38067, 0.42924, 0.47656, 0.52343, 0.57060, 0.61889, 0.66933, 0.72347, 0.78414, 0.85840, 1.00000. Free energies and estimates of the standard error were evaluated using the MBAR method implemented in the edgembar program in the FE-ToolKit package.^90^

### Classical molecular dynamics simulations

All simulations were performed using the AMBER24^161^ software suite. Classical molecular dynamics simulations were performed utilizing GPU accelerated MD.^143^ All energy minimizations were performed with the conjugate gradient method. Temperatures were regulated with a Langevin thermostat^162^ with a collision frequency of 5 ps*^−^*^1^, and pressures were controlled with the Berendsen barostat^163^ during initial equilibration and Monte Carlo barostat^164^ for production simulations. The SHAKE algorithm^165^ was used to fix bonds involving hydrogen. Electrostatics were calculated with the particle mesh Ewald method^166,167^ using a 1 ^Å3^ reciprocal space grid and 12 Å real space cutoffs.^168^ Hydrogen mass repartitioning^169^ (HMR) was used during production MD simulations along with 4 fs time-steps. Simulations were performed in the NPT ensemble with frames collected every 20 ps for statistical analyses.

Starting from the solvent equilibrated structures, first, a 200 ns simulation was run. All systems were tested for RMSD convergence that was reached within the first 100 ns, therefore the first 100 ns were excluded from the analysis. Analysis of RMSD can be found in Figures S20-S22 of the Supporting Information. Independent trials were initiated from structures at time stamps 25, 50, 75, and 100 ns. Each were run for 200 ns, using only the last 100 ns in analysis. This yielded a total of 0.5 *µ*s of production simulation for analysis across 5 trials.

### Catalytic fitness measurements

Catalytic fitness scores were computed to assess how often a given catalytic strategy is satisfied during production MD simulations. Heuristic measurements have been established previously to quantify ideal distances and angles for the four catalytic strategies.^104,170,171^ In each case, the score is computed as

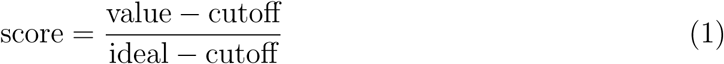

such that the result is 0 if the value falls below the cutoff, 1 if it meets or exceeds the ideal value, and scales from 0 to 1 between the cutoff and ideal values. Metrics with multiple contributions where combined as follows:

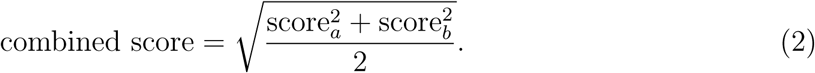

*α* catalysis is a metric that combines inline attack angle (ideal=170°, cutoff=140°) and inline attack distance (ideal=3.1, cutoff=3.5 Å). *β* catalysis combines the distances between the scissile phosphate NBOs to Mg^2+^ (ideal=2.0, cutoff=2.5 Å) or Na^+^ (ideal=2.0, cut-off=3.0 Å). If the metal is Na^+^, the score contribution is scaled by 0.5 to account for less electrostatic stabilization relative to Mg^2+^. *γ* and *δ* catalysis are combined metrics involving Mg^2+^ coordination and hydrogen bonds that vary based on the mechanistic scenario being simulated, a full list of which is available in Table S4.

### Combined quantum mechanical/molecular + machine learning mechanical simulations

Due to the size of the THr active site that needs to be treated quantum mechanically, including catalytic metal ions and their coordination environment, enumerating the reaction pathways for different mechanistic scenarios at the *ab initio* QM level can be prohibitively expensive.^172^ Semi-empirical quantum methods offer much better computational efficiency but are limited in their accuracy due in part to the minimal valence basis set approach, especially for locally anionic regions of the system.^173^ To address this challenge, we train a QM/MM+ΔMLP using the range corrected framework of Zeng et al^89,112^ to correct QM-QM and QM-MM interactions within a 6 Å cutoff. We use the MACE^174,175^ graph neural network as the machine learning potential, which has been enabled by the new DeePMD-GNN plugin^89^ to DeePMD-kit^176^ and recently demonstrated to be transferable for similar chemical reactions in aqueous solution.^90^ All QM/MM+ΔMLP calculations were conducted using interoperable software infrastructure for free energy simulations in AMBER.^144,145^ QM/MM simulations were performed in the canonical ensemble with a 1 fs time-step.

Hydrogens in the QM region were not treated with SHAKE. The boundary between the QM and MM region was treated with the hydrogen link-atom approach.^177,178^ Electrostatics were handled with an Ewald method that uses Mulliken charges,^179^ or ambient potential composite Ewald method^180^ for semi-empirical and *ab initio* theories, respectively, both using 10 Å real space cutoffs.

QM/MM calculations were performed with three separate QM Hamiltonians, *ab initio* PBE0/6-31G*,^181^ DFTB3/3ob tight binding model (using the “3ob” parameter set),^182^ and with a machine learning potential correction, QM/MM+ΔMLP, developed in this study and described separately.

### Mechanistic exploration with string simulations

Mechanisms were studied by quantum mechanical and molecular mechanical (QM/MM) calculations, where the reactive regions of the ribozyme were described by quantum theory. QM regions for both Step 1 and Step 2 reactions included the splice site phosphate, leaving group sugar, nucleophile sugar, 4 surrounding phosphates involved in metal ion coordination (residues C208, C262, A304, and A306), metal ions at A and C (and when present, B) sites, and coordinating waters. In systems where the mechanism for the activation of the nucleophile was explored, a QM hydroxide was also included, otherwise it was a QM water molecule and the nucleophile was deprotonated. The total number of QM atoms was 77 or 90 in the absence or presence of Mg(B), respectively. Mechanisms studied and their corresponding components are summarized in Table S6, with the QM region depicted in Figure S13.

The cleavage and exon ligation steps were both studied by tracking 1 to 3 reaction coordinates that define the mechanism of interest, with phosphoryl transfer coordinate (*ξ*_PT_) defining the splicing in all of them. In addition, a nucleophile activation coordinate (*ξ*_Nu_*_∗_*) and/or leaving group protonation coordinate (*ξ*_LG_*_∗_*) are included for various mechanisms.

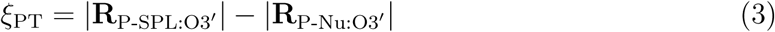

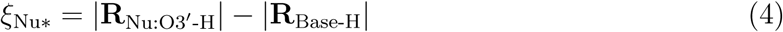

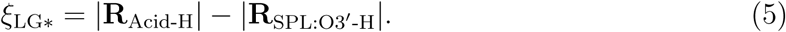

The attacking and leaving O3*^′^* are denoted as Nu for nucleophile and SPL for splice site, respectively, which are in Step 1: Nu=G_exo_ & SPL=U(−1) and in Step 2: Nu=U(−1) & SPL=G*_ω_*. The identities of acids and bases studied in different mechanisms are summarized in Table S6.

Mechanisms involving *ξ*_Nu_*_∗_* and/or *ξ*_LG_*_∗_* (i.e. multiple reaction coordinates), were studied using the surface accelerated string method^91^ included in the FE-ToolKit package.^90^ The initial guess for the minimum free energy path was a straight line connecting the approximate reactant and product states characterized by 32 umbrella windows. A force constant of 100 kcal·mol*^−^*^1Å^*^−^*^2^ was used for the reaction coordinate(s). In each case, *ξ*_PT_ spanned -2.0 Å to 2.0 Å. When present, the reactant/product values of *ξ*_Nu_*_∗_* and/or *ξ*_LG_*_∗_* were determined from MD simulations, with the other end state defined as the negative of that value.

Using QM/MM+ΔMLP as the quantum model enabled sampling of 1 ps per umbrella window per string iteration. Simulations were performed in the canonical ensemble at 298 K using a 1 fs time-step, collecting values of the reaction coordinate(s) every 20 steps. At each iteration, after the sampling, a maximum of 300 “synthetic iterations” were performed optimizing a path with 50 control points and 400 spline points along the current free energy surface to update the string. The free energy surface was approximated by a weighted average of nearby bin values using 4*^th^* order B-splines. All pathways reached the convergence criteria recommended in ref 90 by 50 iterations, and production simulations were performed for 4 independent trials of 25 ps on the resulting MFEP. Standard errors of the free energies were computed by propagation of bootstrap errors.

Intrinsic rate constants were estimated according to the transition state theory,

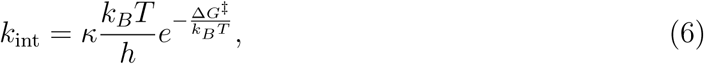

where *h* is Planck’s constant, *k_B_*is the Boltzmann constant, *T* = 298 K, *κ* is the transmission coefficient assumed to be 1, and Δ*G^‡^* is the activation energy obtained from the MFEP predicted with string simulations.

### QM/MM+ΔMLP model training

We trained a machine learning potential correction (ΔMLP) model correcting a low level semi-empirical QM model to a high level *ab initio* reference, with the goal of producing reaction profiles with *ab initio* accuracy at computational costs similar to semi empirical methods. A range-corrected^112^ ΔMLP was trained to correct DFTB3/3ob free energy profiles to the PBE0/6-31G* level based on a MACE^174^ graph neural network potential. Training data was generated by obtaining DFTB3 MFEPs for each reaction. On these paths, 500 structures per window were used to obtain DFTB3 and PBE0 energies and forces through single point calculations, yielding a dataset of 208,000 structures combined across 13 mechanisms studied. Training was performed using the DP-GEN simplify workflow^183^ in DeePMD-Kit version 3.^176^ The hyperparameters used to train the MACE model were consistent with the MACE-OFF medium model.^184^

In short, graph edges include all atomic pairs within a 6 Å cutoff, with edge features expanded in 3^rd^ order spherical harmonics and 8 radial Bessel functions, smoothly decaying to zero at the cutoff via a 5^th^ order polynomial envelope. These features are passed through a 3 layer multilayer perceptron with 64 neurons per layer. Equivariant message passing uses first order spherical harmonics with 128 embedding channels over 2 interaction layers. A correlation order of 3 was used to yield 4-body features. Final atomic energies were obtained from a 16 channel perceptron with SiLU activation.

The training procedure performed by the DP-GEN simplify program is a pool-based active learning approach. A subset of structures from the full dataset (pool) are iteratively used to train 4 stochastic models. After an iteration of training, the models are evaluated on the remaining data set to compute the deviation in forces predicted by the 4 models, with high deviation corresponding to high uncertainty. The model deviation is defined as:

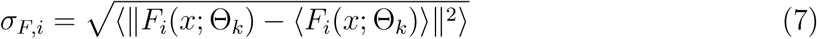

where *F_i_* is the atomic force for a given data frame *x*, and Θ is the set of parameters defining the model *k*. Structures with model deviation falling below a 0.08 eV/Å (1.84 kcal·mol*^−^*^1Å^*^−^*^1^) cutoff are considered accurate and removed from the dataset, such that the dataset is “simplified” to focus training on statistically independent samples. The initial training iteration used 20,000 samples, with subsequent iterations adding 10,000 new samples. The neural network parameters were updated from 100,000 steps of Adam stochastic optimization^185^ with an initial learning rate of 0.001 that decayed exponentially to 5×10*^−^*^8^. The loss function contains contributions from energies and forces and the prefactor associated with each property has a dependence on the learning rate. The initial and final prefactors of the energy term are 0.02 eV*^−^*^2^ and 1.0 eV*^−^*^2^, respectively, and the initial and final prefactors of the force terms are 1000 eV*^−^*^2Å2^ and 1.0 eV*^−^*^2Å2^. A detailed description of the loss function can be found in ref. 186.

### Predictions of p*K*_a_s from quantum chemical calculations

Inspired by the up-shifted p*K*_a_ at the N1 position of 7cdiAP relative to diAP experimentally,^139^ we sought to estimate the p*K*_a_s of 3cdiAP and 3,7cdiAP to gauge whether such compounds, if synthesized, could exhibit further up-shift to stabilize a wobble pair with C with full occupancy. Given that p*K*_a_s for these compounds were not found in the literature, we employed *ab initio* quantum chemical calculations to make estimates. The free energy of protonation for species A is given as Eq. 8, where the standard free energy of the proton in water (−270.297 kcal·mol*^−^*^1^) was used as in other work.^187^

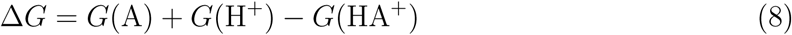

The corresponding calculated p*K*_a_ values were determined as:^187^

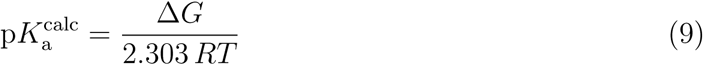

where R is the gas constant and T is temperature (298 K).

Previous studies have shown that reliable p*K*_a_ values can be obtained using methyl-substituted nucleobase models, where the ribose moiety is replaced by a methyl group.^187^ This strategy was therefore employed in the present work to reduce computational cost. Estimating the p*K*_a_ directly from quantum calculations can be a challenge, so we first constructed a regression model using 9 modified nucleobases (Tables S8) with experimentally known p*K*_a_s, inspired by ref 187 and previously used in ref 188, to benchmark the relationship between experimental and quantum chemically predicted p*K*_a_s. From this regression model, we predict the experimentally unknown p*K*_a_s as follows:

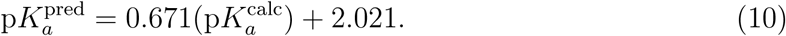

Electronic structure calculations were performed using Gaussian 16.^189^ Each molecule in the protonated and deprotonated state were subjected to geometry optimization and frequency calculations at the M06-2X/aug-cc-pVTZ level of theory^190,191^ with the SMD implicit solvation model.^192^ Tight convergence criteria and an ultrafine integration grid were used throughout. Additional details regarding the compounds used and validation tests are given in the Supporting Information.

## Source data

Additional source data for PBSA, 1D-RISM, 3D-RISM, MD simulations, ABFE simulations, QM/MM+ΔMLP training, and QM/MM+ΔMLP simulations are available online through Zenodo: 10.5281/zenodo.21221644. Source data includes AMBER mdin files, non-standard residue parameters, MD trajectories, free energy analysis, MACE model files and training inputs, and free energy profile data.

## Supporting Information

The Supporting Information contains Figures S1-S22, Tables S1-11, and supporting discussion. These include additional overview of the available cryo-EM structures, analysis of 3D-RISM calculations, analysis of alchemical free energy simulations, analysis of MD simulations, QM setup details, DFTB3 level free energy profiles, machine learning potential validation, additional analysis of free energy profiles, and details of QM p*K*_a_ predictions.

## Author Information

### Contributions

E.M. designed, performed, and analyzed the simulations and 3D-RISM calculations; trained machine learning models; and wrote the original draft of the manuscript. Ş.E. and T.J.G. contributed to the development of free energy sampling and analysis methods. Ş.E. provided non-standard residue parameters. T.J.G. developed the QM/MM+ΔMLP architecture in AMBER. Y.D. contributed computational p*K*_a_ predictions. G.P.B. developed AFE workflow infrastructure. A.W.G. assisted with machine learning training and data collection. Ş.E., T.J.G., J.A.P., and D.M.Y. provided critical evaluation and data interpretation. D.M.Y. conceived the project and supervised all aspects of the research. All authors reviewed, edited, and approved the final version of the manuscript.

### Ethics declarations

The authors declare no competing interests.

## Acknowledgments

Computational resources were provided by the Office of Advanced Research Computing (OARC) at Rutgers, The State University of New Jersey; the San Diego Supercomputer Center (SDSC) at the University of California San Diego (supercomputer Cosmos supported by National Science Foundation Grant No. 2404323); the Advanced Cyberinfrastructure Coordination Ecosystem: Services & Support (ACCESS) program, which is supported by National Science Foundation grants #2138259, #2138286, #2138307, #2137603, and #2138296 (supercomputer Expanse at SDSC through allocation CHE190067); the Texas Advanced Computing Center (TACC) at the University of Texas at Austin, URL: http://www.tacc.utexas.edu (supercomputer Frontera through allocation CHE20002).

