## Supporting Information for "A transient third metal ion shapes the catalytic cycle of a group I intron ribozyme"

### Overview of the splicing reaction and available cryo-EM structures

In this section we provide a brief orientation regarding the splicing reaction catalyzed by THr, and how it appears in the cryo-EM structures[S1–S3] used to initiate simulations in this work. The L-16 S*ca*I *Tetrahymena* ribozyme performs splicing in trans, meaning the 5′ exon-intron oligonucleotide is not explicitly bonded to the ribozyme. This difference is highlighted in Figure S1. The structure of the ribozyme, including its helical domains, are represented in greater detail in Figure S2. Finally, we provide additional views of the electrostatic potential surfaces computed on the cyro-EM structures that depict the substrate surfaces at the active site.

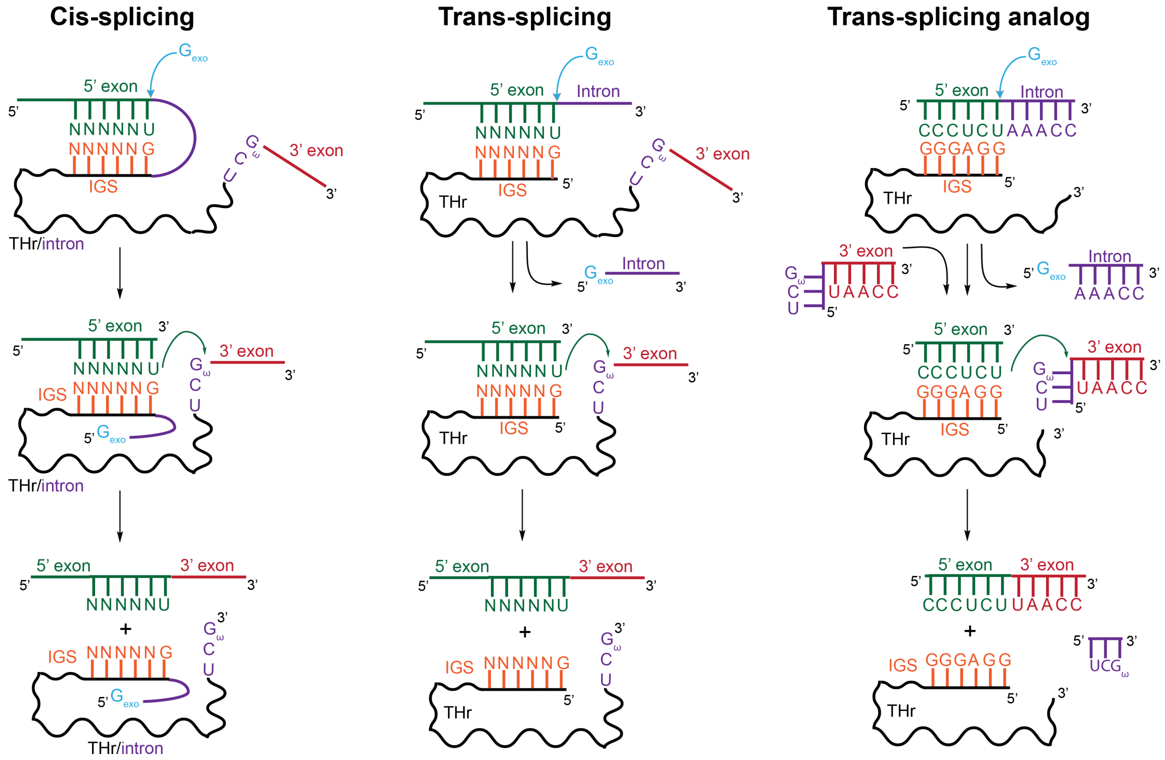


Figure S1. Schematic overview of the general mechanisms of cis and trans-splicing catalyzed by THr. The trans-splicing analog represents the sequences and fragments present in the structures simulated in this study. IGS refers to the internal guide sequence of the ribozyme. Residues labeled “N” could be tuned for splicing reactions involving desired nucleotides.

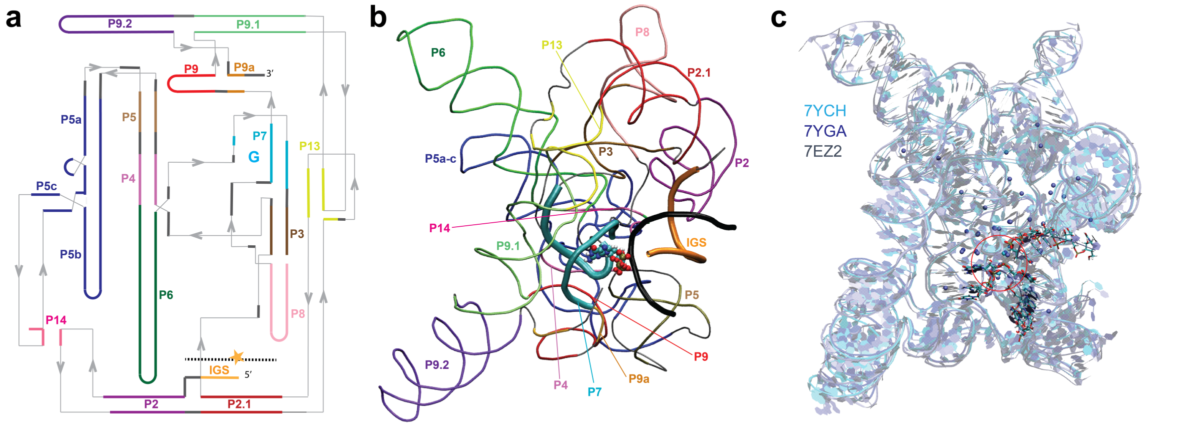


Figure S2. Cryo-EM structures of THr used to initiate simulations. **(a)** Overview of the THr domains and connectivity. The G binding site is indicated by a cyan colored "G" bound by the P7 domain. The substrate is indicated by a black dashed line, with the splice site denoted by a star. **(b)** Representation of the 7YCH structure prior to cleavage colored by domain. The P7 domain (cyan), internal guide sequence (IGS, orange), and substrate (black) are emphasized, and GTP is represented atomistically. **(c)** Alignment of the pre-catalytic ribozymes from 7YCH (cyan), 7YGA (dark blue), and 7EZ2 (gray) containing resolved $Mg^{2+}$ ions (spheres). The bound RNA strands of 7YGA are represented atomistically, and were not included in the alignment.

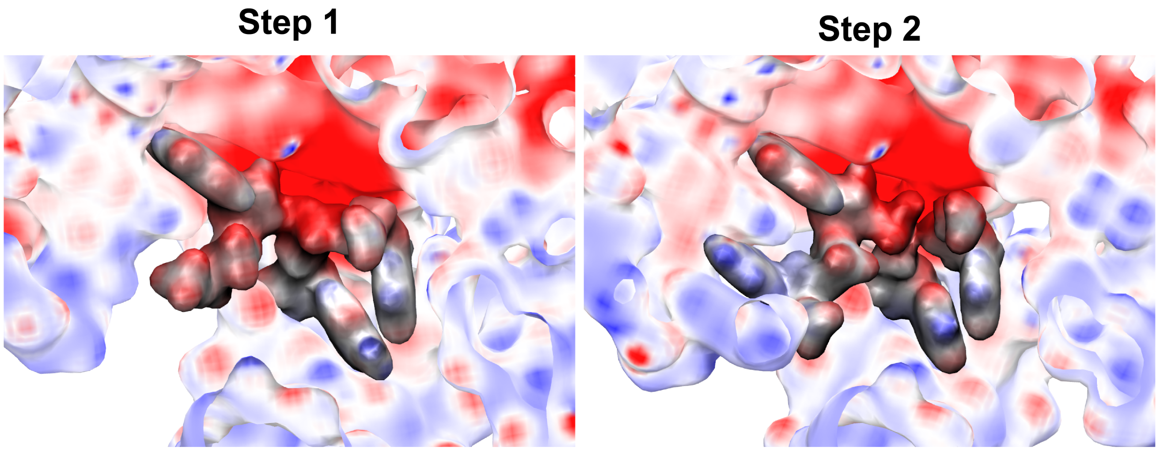


Figure S3. Electrostatic potential surface representations of the active site including substrates of the cleavage (Step 1) and ligation (Step 2) structures from Poisson-Boltzmann Surface Area calculations. Red coloring indicates negative charge, and blue positive charge.

### Extended analysis of 3D-RISM isocountour values

In order to probe for high probability $Mg^{2+}$ binding sites, we used 3 dimensional reference interaction site modeling (3D-RISM) to compute the $Mg^{2+}$ probability density around the ribozyme relative to bulk solution containing 10 mM $Mg^{2+}$, 140 mM $Na^{+}$, and 160 mM $Cl^{-}$. Metal ion binding site coordination spheres are defined in Table S1. In addition, binding modes resolved in the related pre-cleavage structure of the *Azoarcus* group I intron[S4] (Azo) are provided. The values reported in Table S2 represent the ratio of the probability density of $Mg^{2+}$ occurrence at the 5 sites closest to the active site relative to the bulk. An accompanying depiction of the 3D-RISM densities with the surrounding ribozyme environment is provided is Figure S4, with the sites labeled in Figure S5.

Table S1. Active site phosphate $Mg^{2+}$ coordination environments simulated by MD based on cryo-EM structures, phosphorothioate substitutions, and 3D-RISM

| $Mg^{2+}$ site | THr Cryo-EM | Ligands | Azo Cryo-EM | Ligands |
|---|---|---|---|---|
| A | Yes | SPL(pro-$S_P$), C208(pro-$S_P$), A304(pro-$S_P$), A306(pro-$S_P$) | Yes | C87(pro-$S_P$), G169(pro-$S_P$), A171(pro-$S_P$) |
| B | No | SPL(pro-$R_P$/pro-$S_P$) | No | - |
| C | No | SPL(pro-$S_P$), C262(pro-$S_P$), A306(pro-$R_P$) | Yes | G127(pro-$S_P$), A171(pro-$R_P$) |
| D | Yes | C262(pro-$R_P$), A207(pro-$S_P$) | Yes | A86(pro-$S_P$) |
| E | Yes | U307(pro-$R_P$), A308(pro-$R_P$) | Yes | U172(pro-$R_P$), A173(pro-$R_P$) |

$Mg^{2+}$ binding sites correspond to labels in Figure S5. The cryo-EM column indicates whether sites were occupied in the cryo-EM structures used to initiate simulations. SPL refers to the scissile phosphate at the splice site. Azo ligands are from the cryo-EM structure 9VNE.

Table S2. Maximum 3D-RISM isocontour values and percentages of maximum for 5 conserved $Mg^{2+}$ ions near the active site

| | | $\rho_{\mathrm{site}}/\rho_{\mathrm{bulk}}$ | | | | | |
|---|---|---|---|---|---|---|---|
| Step | Construct | Mg(A) | Mg(C) | Mg(B) | Mg(D) | Mg(E) | max |
| 1 | GTP, SS | 94 (99%) | 83 (87%) | 65 (68%) | 76 (80%) | 68 (71%) | 95 |
| 1 | GTP, AP | 109 (79%) | 136 (99%) | 68 (49%) | 83 (60%) | 90 (66%) | 137 |
| 1 | UCG, AP | 126 (89%) | 141 (99%) | 77 (54%) | 84 (59%) | 86 (61%) | 142 |
| 1 | G, AP | 122 (96%) | 122 (96%) | 63 (49%) | 82 (64%) | 68 (53%) | 128 |
| 2 | SS | 109 (99%) | 91 (82%) | 80 (72%) | 79 (71%) | 78 (71%) | 111 |
| 2 | AP | 145 (99%) | 101 (69%) | 74 (51%) | 64 (44%) | 82 (56%) | 147 |

SS indicates the standard state with the nucleophile protonated, and AP the activated precursor state with nucleophile deprotonated. Metal ion sites are labeled in Figure S5. Percentages of the maximum are given in parentheses.

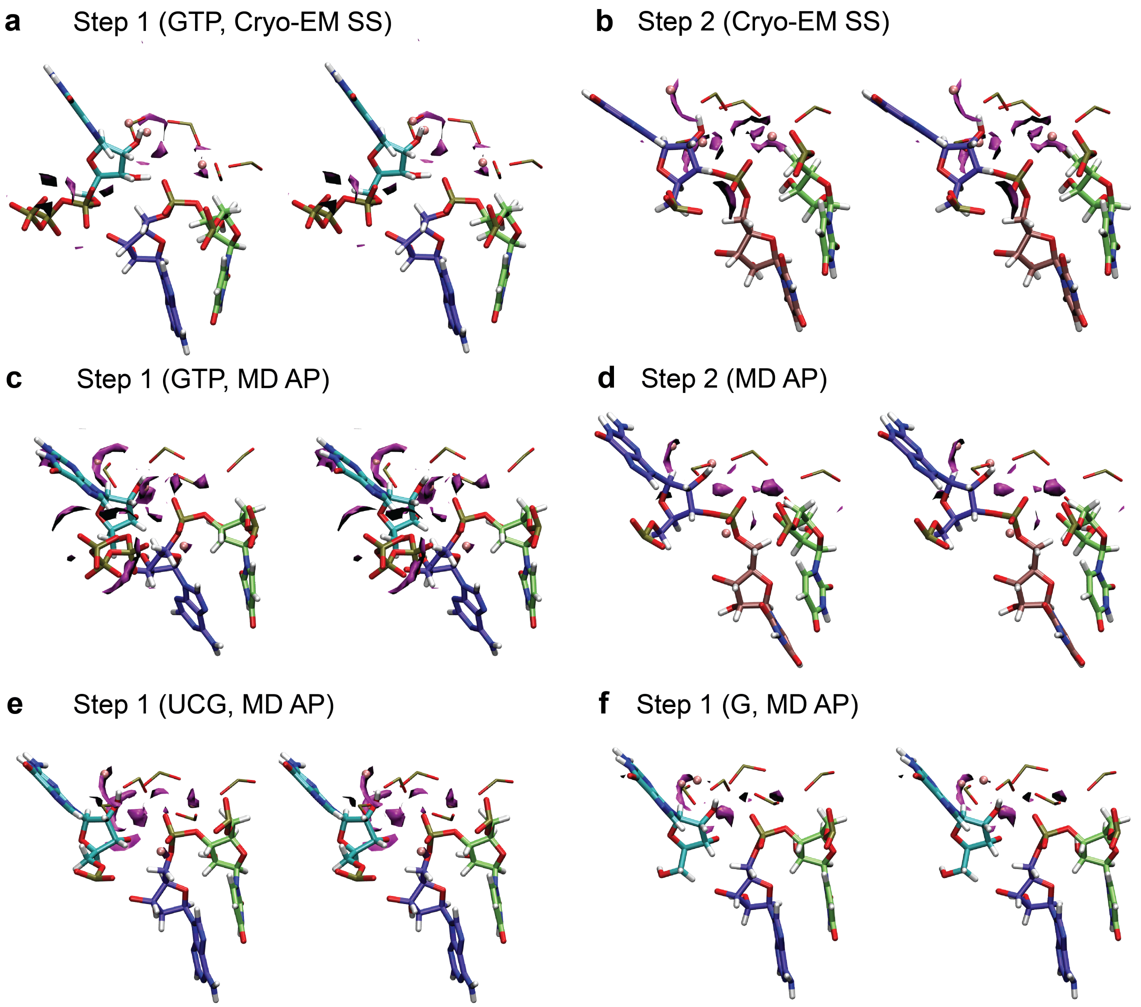


Figure S4. Active site $Mg^{2+}$ ion isodensity contours predicted by 3D-RISM for **(a)** Step 1 (cleavage) and **(b)** Step 2 (ligation) in the standard state (SS) from cryo-EM with nucleophile protontated. 3D-RISM isodensity contours for **(c)** Step 1 and **(d)** Step 2 in the nucleophile deprotonated, activated precursor (AP) state from MD simulations. 3D-RISM isodensity contours for the AP state of Step 1 where $G_{exo}$ is **(e)** UCG and **(f)** guanosine (G). $Mg^{2+}$ (magenta) binding densities are calculated from 3D-RISM and depicted at the 66$^{th}$ contour level. Structures are depicted in parallel stereo.

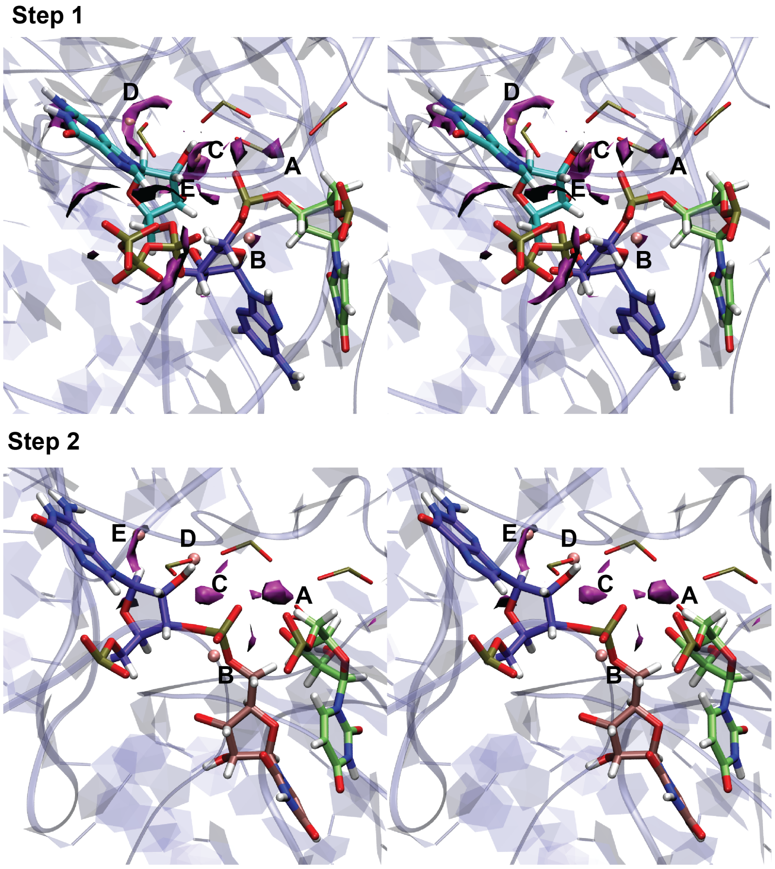


Figure S5. Stereo images of the $Mg^{2+}$ binding sites A through E labeled in the context of the first and second steps of splicing in the activated precursor state.

### Extended analysis of $Mg^{2+}$ ion binding free energies

Absolute binding free energy (ABFE) simulations were used to estimate the relative binding affinities between active site metal ions. Expanding on 3D-RISM, this approach accounts for dynamics to provide a more rigorous, quantitative view of the differences in affinities of sites between Step 1 and 2. To contextualize the free energies, Table 1 reports a $\Delta\Delta G_b$ between the ribozyme environment and a model GpG dinucleotide environment where the $Mg^{2+}$ ion coordinates a single phosphate, as shown in Figure S6. In this way, it is clear that, even when 5 $Mg^{2+}$ ions are present in close proximity, the highly negatively charged nature of the active site makes each binding site more favorable than in the dinucleotide.

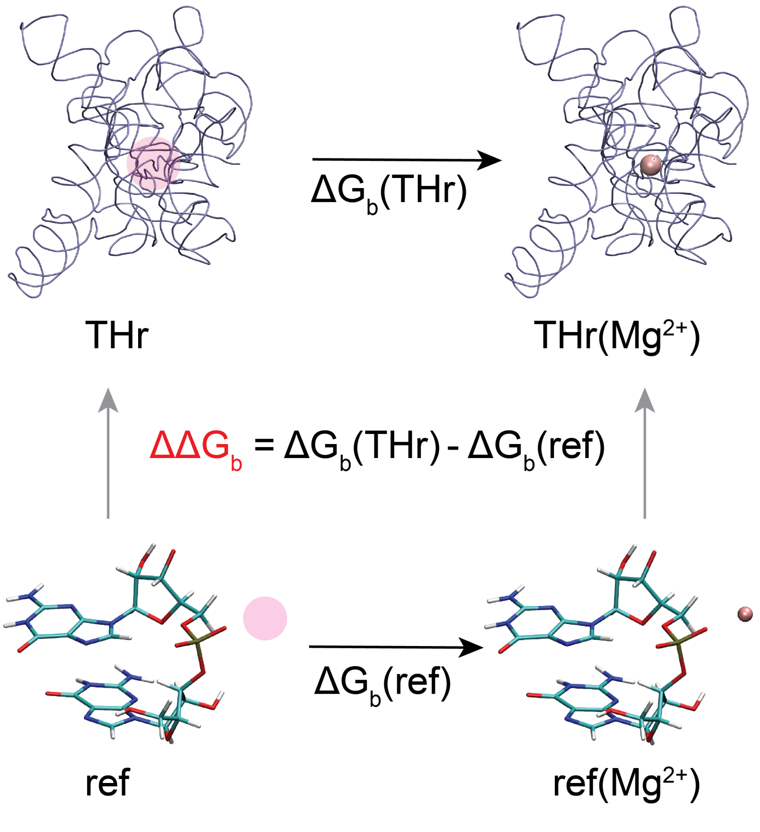


Figure S6. Thermodynamic cycle used to estimate relative binding free energies ($\Delta\Delta G_b$) of active site $Mg^{2+}$ ions. The free energy is reported as $\Delta\Delta G_b$ between the ion being annihilated in the ribozyme environment (THr) and in an environment directly coordinating a single phosphate in the dinucleotide GpG (ref), where $\Delta G_b$ values are obtained from absolute binding free energy simulations. A translucent sphere indicates the $Mg^{2+}$ ion is in the dummy state.

As shown in Table S3, the binding free energies generally increased upon formation of the activated precursor when the nucleophile interacts with that site. Mg(A) in the AP state of Step 2 was an exception. This was due to a $Na^+$ ion from solution entering the A binding site while Mg(A) was in the dummy state, lowering the free energy compared to other simulations where $Na^+$ did not infiltrate the binding sites. There is also a marked increase in the error estimates in the free energies when very high affinity sites are eliminated due to structural adjustments in response to the electrostatic repulsion of phosphates, and differences in the solvent environment between trials.

Table S3. Absolute binding free energies of active site $Mg^{2+}$ ions computed from alchemical free energy simulations

| | | | ΔG (kcal·mol$^{-1}$) | | | | |
|---|---|---|---|---|---|---|---|
| $Mg^{2+}$ | Step | State | tr1 | tr2 | tr3 | mean | $\sigma$ |
| Soln | - | - | -369.2 | -369.4 | -369.6 | -369.4 | 0.2 |
| GpG | - | - | -436.6 | -436.4 | -436.2 | -436.4 | 0.1 |
| A | 1 | SS | -479.0 | -476.9 | -478.1 | -478.0 | 0.6 |
| | | AP | -483.8 | -485.8 | -484.3 | -484.6 | 0.6 |
| B | | SS | -440.3 | -440.2 | -442.4 | -441.0 | 0.7 |
| | | AP | -444.0 | -443.3 | -444.3 | -443.9 | 0.3 |
| C | | SS | -455.7 | -454.5 | -455.4 | -455.2 | 0.4 |
| | | AP | -491.4 | -483.7 | -484.5 | -486.5 | 2.5 |
| D | | SS | -454.2 | -455.6 | -451.3 | -453.7 | 1.3 |
| | | AP | -448.7 | -451.5 | -453.7 | -451.3 | 1.5 |
| E | | SS | -453.8 | -452.6 | -457.1 | -454.5 | 1.4 |
| | | AP | -454.2 | -451.7 | -452.1 | -452.7 | 0.8 |
| A | 2 | SS | -489.7 | -485.6 | -482.0 | -485.8 | 2.2 |
| | | AP* | -473.5 | -479.7 | -473.4 | -475.6 | 2.1 |
| B | | SS | -451.2 | -447.5 | -449.6 | -449.4 | 1.1 |
| | | AP | -447.7 | -449.4 | -447.7 | -448.3 | 0.6 |
| C | | SS | -460.6 | -457.1 | -467.9 | -461.9 | 3.2 |
| | | AP | -464.8 | -464.8 | -474.6 | -468.1 | 3.3 |
| D | | SS | -446.4 | -445.0 | -447.7 | -446.4 | 0.9 |
| | | AP | -450.7 | -448.7 | -450.5 | -450.0 | 0.6 |
| E | | SS | -451.0 | -451.6 | -448.7 | -450.4 | 0.9 |
| | | AP | -459.0 | -453.6 | -454.9 | -455.8 | 1.7 |

All simulations for Step 1 were performed with $GTP_{exo}$. Free energies were computed using MBAR, and $\sigma$ is the uncertainty computed from the circular moving block bootstrap procedure described in ref S5. A "*" indicates that a $Na^+$ ion entered the Mg(A) binding site upon annihilation. GpG is a dinucleotide where the $Mg^{2+}$ ion coordinates a single phosphate, and soln is a $Mg^{2+}$ ion in water.

### Detailed analysis of catalytic fitness

Catalytic fitness scoring was used to assess the requirements for a catalytically poised active state ensemble. In general, we sought to understand whether metal ion binding at site B was necessary to establish high catalytic fitness, and how this requirement may change from Step 1 to Step 2, or in the presence of different $G_{exo}$ species. The metrics used to quantify catalytic fitness are given in Table S4. Table S5 provides average fitness scores corresponding to Figure 3 in the main text. In Figures S7-S9 we provide a more detailed view of the per-system catalytic fitness.

Table S4. Ideal and cutoff angles and distances used to compute catalytic fitness scores

| | | °/Å | |
|---|---|---|---|
| Strategy | Angle/Distance | Ideal | Cutoff |
| $\alpha$ | Nuc-P-LG | 170 | 140 |
| | Nuc-P | 3.1 | 3.5 |
| $\beta$ | NBO-$Mg^{2+}$ | 2.0 | 2.5 |
| | NBO-$Na^{+}$ | 2.0 | 3.0 |
| $\gamma$ | H-Nuc | 1.7 | 2.5 |
| $\delta$ | H-LG | 1.7 | 2.5 |

Nuc refers to the nucleophile O3′, P to the phosphorous of the scissile phosphate, and LG to the O3′ leaving group. H is the hydrogen from hydrogen bond donors including water or O2′.

As shown in Figure S7, the first step maintains high catalytic fitness in the presence and absence of Mg(B) in both the SS and AP states. In these simulations, GTP is bound at the $G_{exo}$ position, and Mg(B) is bound directly coordinated to the pro-$R_P$ NBO and 5 water molecules, the position observed from 3D-RISM. This binding mode results in indirect coordination to the leaving group and the U(-1) phosphate. This leaving group hydrogen bond also contributes to $\delta$ catalysis. Due to the $G_{exo}$ sugar pucker, the nucleophillic O3′ is not able to form a hydrogen bond with water on the solvent facing side of the active site, therefore $\gamma$ catalysis is not impacted by Mg(B) in this binding mode. The triphosphate group of GTP is rotatable, and may oscillate between direct and indirect coordination with Mg(B).

Table S5. Average catalytic fitness scores from MD simulations of THr under various active site conditions

| System | | | Score | | | | |
|---|---|---|---|---|---|---|---|
| Step | $G_{exo}$ | Mg(B) | $\alpha$ | $\beta$ | $\gamma$ | $\delta$ | mean |
| 1 | GTP | - | 0.64 | 0.48 | 0.76 | 0.71 | 0.70 |
| 1 | GTP | NBO | 0.61 | 0.78 | 0.69 | 0.71 | 0.77 |
| 2 | - | - | 0.50 | 0.46 | 0.00 | 0.59 | 0.63 |
| 2 | - | NBO | 0.72 | 0.78 | 0.60 | 0.41 | 0.72 |
| 2 | - | NBO,Nuc | 0.49 | 0.70 | 0.53 | 0.45 | 0.60 |
| 2 | - | NBO,LG | 0.00 | 0.78 | 0.59 | 0.00 | 0.44 |
| 1 | G | - | 0.68 | 0.48 | - | 0.69 | 0.71 |
| 1 | UCG | - | 0.67 | 0.48 | 0.74 | 0.69 | 0.71 |
| 1 | UCG | NBO | 0.77 | 0.80 | 0.54 | 0.69 | 0.81 |

Scores for $\gamma$ fitness are taken the SS, and remaining scores are taken from the AP. Mg(B) is either not present (-), directly coordinating the NBO, or directly coordinating the NBO along with the nucleophile (Nuc) or leaving group (LG). Other $Mg^{2+}$ binding modes are as indicated in Figure 3. $\alpha$, $\beta$, $\gamma$, and $\delta$ catalysis correspond to the interactions depicted in Figure 3, and scores were computed as reported in Equations 1 and 2.

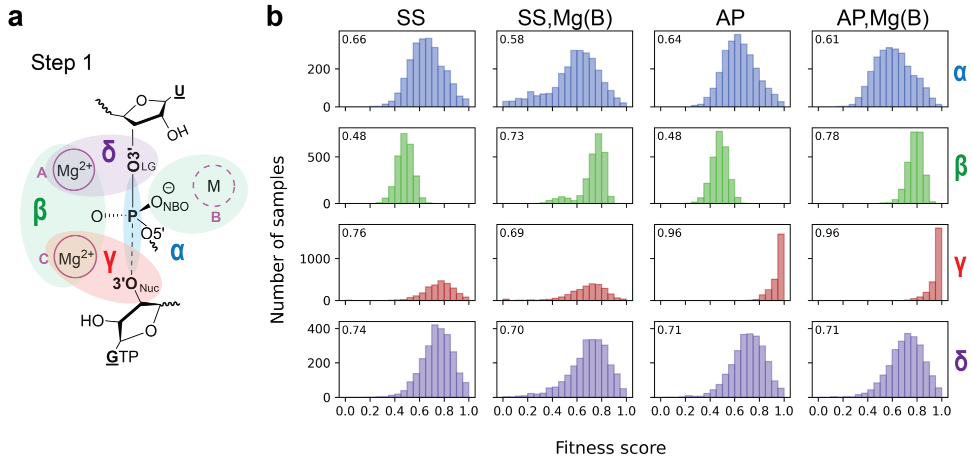


Figure S7. Catalytic fitness scores from MD simulations of the first step of splicing. **(a)** Schematic representation of the catalytic strategies used in Step 1, with $Mg^{2+}$ coordinating the pro-$R_P$ NBO. **(b)** Distributions of catalytic fitness for the standard state (SS) and activated precursor (AP) with and without Mg(B). Average scores are given in the top left corner.

Next we looked at the impact of different $G_{exo}$ species on catalytic fitness. As shown in Figure S8, guanosine (G) and UCG exhibit high catalytic fitness, on par with GTP. However, when Mg(B) is added to the UCG system, it becomes the most catalytically fit species. In this case, Mg(B) is directly coordinated to the pro-$R_P$ NBO, and indirectly coordinated to the leaving group, U(-1) phosphate, and G(+1) phosphate. The presence of this rigid phosphate upstream of the cleavage site appears to create a well defined Mg(B) site that improves catalytic fitness.

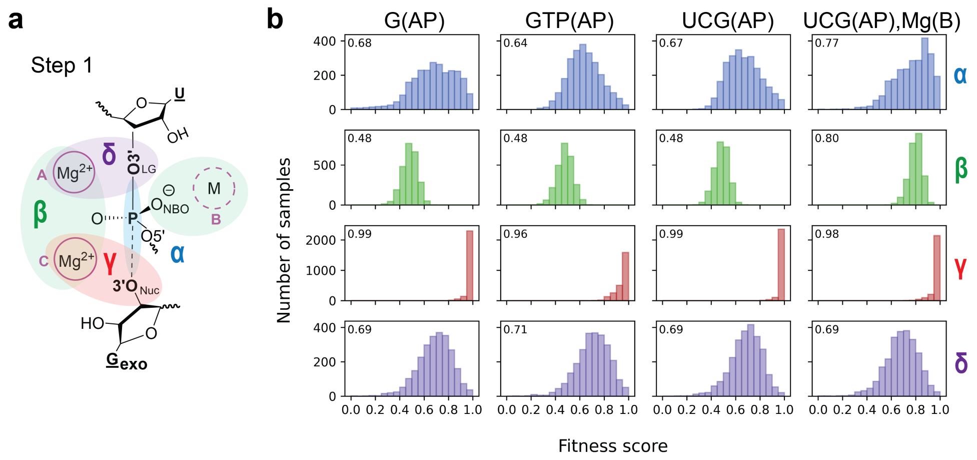


Figure S8. Catalytic fitness scores from MD simulations of the first step of splicing with different $G_{exo}$ moieties. **(a)** Schematic representation of the catalytic strategies used in Step 1, with $Mg^{2+}$ coordinating the pro-$R_P$ NBO. **(b)** Distributions of catalytic fitness for the activated precursor state of Step 1 in the presence of G, GTP, and UCG. Average scores are given in the top left corner.

In the second step of splicing, inline fitness was significantly reduced upon forming the AP state, as shown in Figure S9. This was not observed in Step 1. However, introduction of Mg(B) directly coordinating the pro-$S_P$ NBO rescued this fitness. Based on these results, one may speculate that perhaps Mg(B) binding is more crucial for Step 2 than Step 1. In addition, the SS state of Step 2 lost inline fitness upon introduction of Mg(B) begging the question of whether Mg(B) binding could be transient, and/or coupled with AP formation.

Experiments have suggested that the O3′ and O2′ of $G_{exo}$ in Step 1 may bind distinct $Mg^{2+}$ ions, a theory that initially raised the possibility of a third metal ion.[S6] This binding mode theory has been challenged more recently.[S7] However, the 3D-RISM results for the AP state of Step 2 showed a distribution of density between the pro-$S_P$ NBO and the nucleophile, suggesting that direct coordination of Mg(B) to both the NBO and nucleophile at the same time may be possible. This would be analogous to the mode proposed by Shan et al in Step 1.[S6] If this were in fact the preferred binding mode, it was not immediately clear whether

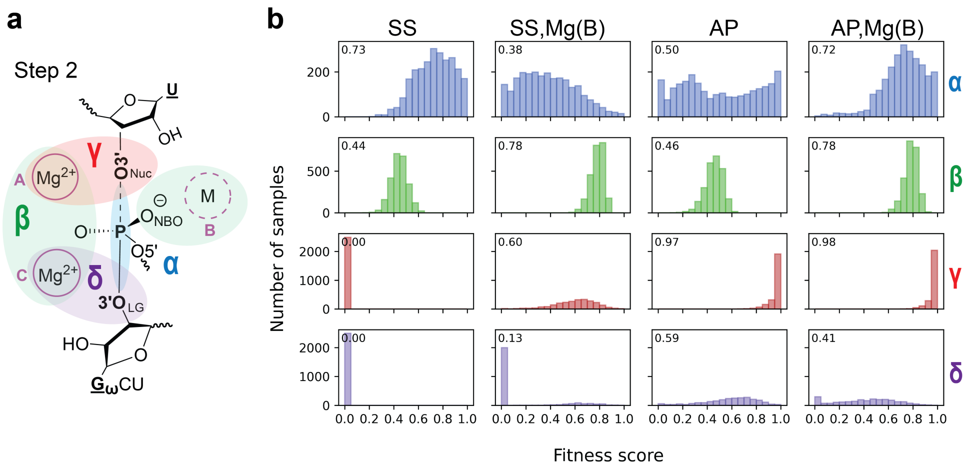


Figure S9. Catalytic fitness scores from MD simulations of the second step of splicing. **(a)** Schematic representation of the catalytic strategies used in Step 2, with $Mg^{2+}$ coordinating the pro-$R_P$ NBO. **(b)** Distributions of catalytic fitness for the standard state (SS) and activated precursor (AP) with and without Mg(B). Average scores are given in the top left corner.

the metal ion would change position and coordinate the nucleophile in both reactions, or hold its position and coordinate the leaving group. Therefore, Mg(B) binding modes with direct coordination to the NBO and either the nucleophile or leaving group were simulated for the first and second step in the AP state. These modes are shown in Figure S10a, with fitness results shown in Figure S10b.

In both Step 1 and Step 2, the leaving group binding mode could not be maintained without restraints. Despite attempts to gradually remove restraints and equilibrate the binding mode, leaving group direct coordination was lost. Therefore, fitness reported for this mode are in the presence of 25 kcal·mol$^{-1}$Å$^{-1}$ restraints maintaining a maximum Mg(B)-leaving group distance of 3.5 Å. For both steps, this mode yielded poor catalytic fitness. Due to the orientation of the nucleophile away from solvent in Step 1, direct coordination led to extended inline distances as the ion came between the nucleophile and electrophile, yielding poor inline fitness and physically blocking attack. This binding mode was less disruptive in Step 2, where the nucleophile faces solvent, but the inline fitness score was significantly worse than when Mg(B) only directly coordinated the NBO.

Finally, we provide a breakdown of $Na^+$ ion accumulation at the solvent exposed NBO,

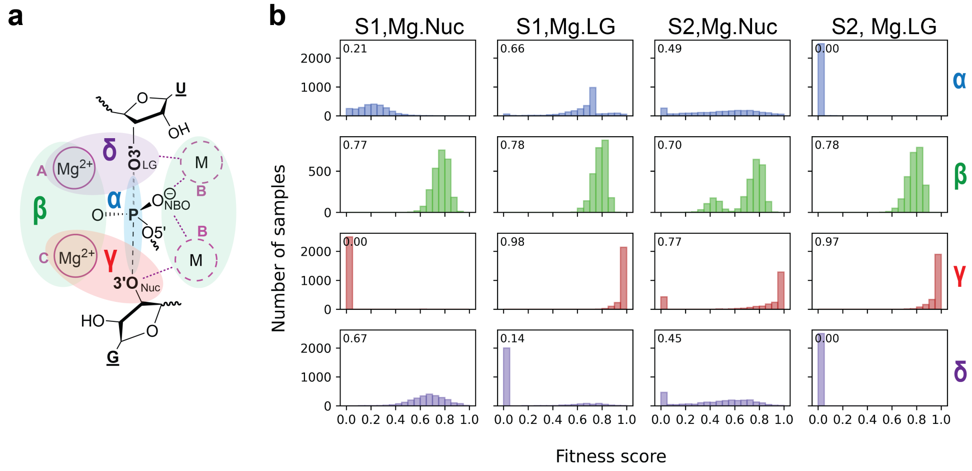


Figure S10. Catalytic fitness scores from MD simulations of the first and second steps of splicing. **(a)** Schematic representation of the catalytic strategies and Mg(B) binding modes tested by MD simulations. **(b)** Distributions of catalytic fitness for the activated precursor state of Steps 1 and 2 with Mg(B) coordinated the NBO as well as either the leaving group (LG) or nucleophile (Nuc). Average scores are given in the top left corner.

which can also contribute to $\beta$ catalysis. It is clear that accumulation in the B site is most prevalent in Step 2, whereas Step 1 accumulation is sensitive to the presense of a phosphate group on $G_{exo}$. In addition, the hydrogen bonding distance of the HO2′ to the phosphates of C262 and U307, with U307 being the primary hydrogen bond partner, is given in Figure S12.

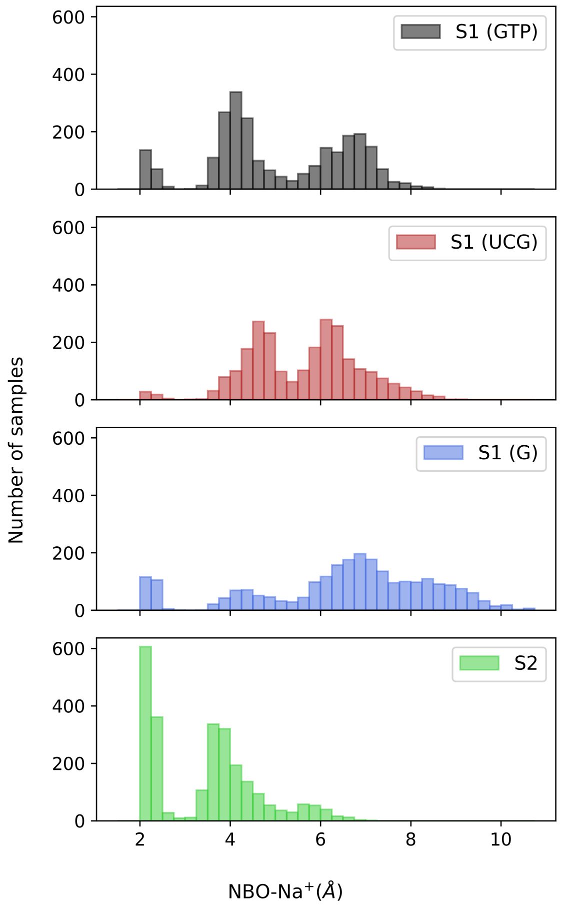


Figure S11. Distance between the solvent exposed NBO and closest $Na^+$ in the AP state of Step 1 with GTP, UCG, and G, and Step 2 from MD simulations.

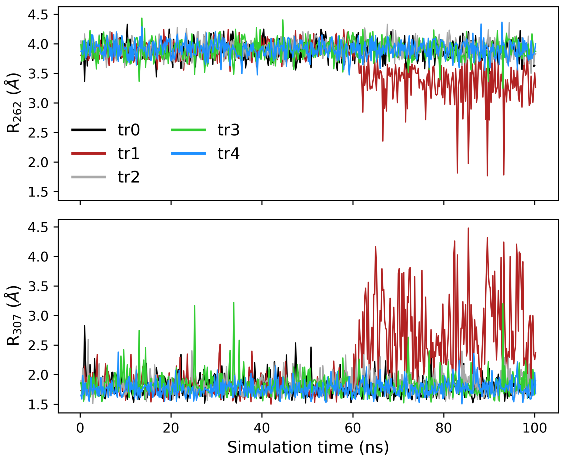


Figure S12. Distances between the HO2′ of G and pro-$R_P$ phosphate oxygens of C262 and U307 for simulations of Step 1 with guanosine bound.

### Detailed description of QM system

A total of 13 mechanisms were probed first by DFTB3 simulations to generate structures for training, then reevaluated with PBE0/6-31G* to obtain high level reference data. Finally, reaction path-finding was performed with QM/MM+ΔMLP simulations. Mechanisms probed involve varying roles played by catalytic components, reaction coordinate dimensionality, and QM region composition. These are summarized in Table S6, with the QM region depicted in Figure S13.

Table S6. Summary of all mechanisms tested by QM/MM simulations

| Nuc | B site | Acid | Base | Ndim | QM charge |
|---|---|---|---|---|---|
| Step 1 | $G_{exo}$ + U-A → $U^*$ + $G_{exo}$-A | | | | |
| GTP | - | - | $OH^-$ | 2d | -2 |
| GTP | - | Wat | $OH^-$ | 3d | -2 |
| GTP | - | LG HO2′ | $OH^-$ | 3d | -2 |
| GTP | - | (A).Wat | $OH^-$ | 3d | -2 |
| GTP | Na | (B).Wat | $OH^-$ | 3d | -1 |
| GTP | Mg | (B).Wat | $OH^-$ | 3d | 0 |
| G- | - | - | - | 1d | -1 |
| UCG- | - | | - | 1d | -2 |
| UCG- | Mg | (B) Wat | - | 2d | 0 |
| Step 2 | $U^*$ + $G_\omega$-U → $G_\omega$ + $U^*$-U | | | | |
| 5′exon U | Mg | Wat | (B).$OH^-$ | 3d | 0 |
| 5′exon U- | -* | - | - | 1d | -1 |
| 5′exon U | Na | - | (B).$OH^-$ | 2d | -1 |
| 5′exon U | Mg | - | (B).$OH^-$ | 2d | 0 |

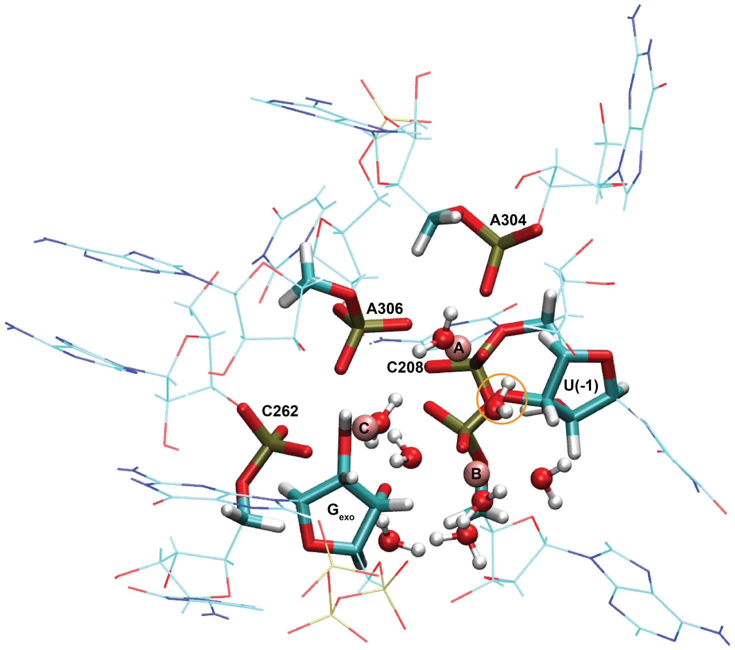


Figure S13. Representative QM region of the Step 1 reaction containing Mg(B). The QM region is represented in licorice and flanking MM residues with sticks. QM regions that lack Mg(B) retain the water circled in orange hydrogen bonding to the leaving group, and Step 2 contains the same QM region as Step 1.

### DFTB3 level free energy profiles

DFTB3 level QM/MM path-finding simulations were conducted to generate training data for the QM/MM+ΔMLP model. In Figure S14 we provide all of the DFTB3 level free energy profiles associated with Figures 4 and 5 in the main text. This is in part to emphasize the importance of using an *ab initio* level QM potential for modeling ribozyme reactions. The free energy barriers obtained at the DFTB3 level gave severe overestimates relative to experiment, suggesting that no mechanistic scenario tested would have been experimentally consistent. In addition, some of the thermodynamic trends do not comport with the *ab initio* results. For example, all of the DFTB3 profiles for Step 1 show deprotonation of the nucleophile by $OH^-$ is thermodynamically unfavorable by ∼12, suggesting the p$K_a$ of O3′ is higher than that of $OH^-$ by ∼ 9 p$K_a$ units. This is physically unreasonable, given the p$K_a$ of O2′ in solution is on the order of 12-13.[S8] This does not even take into consideration that the O3′ directly coordinates a metal ion. Given that DFTB3 uses a minimal basis set approach, the highly negatively charged junctions of nucleic acids are a challenge and should be treated with an appropriate level of theory for comparison with experimental trends.

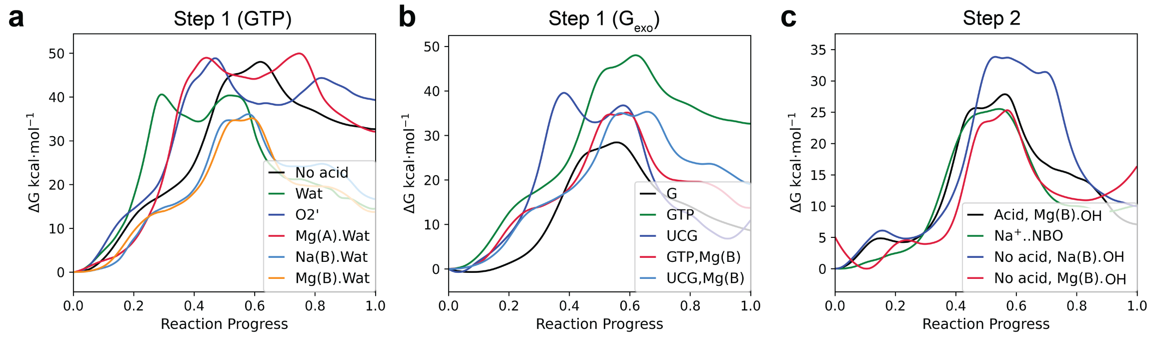


Figure S14. DFTB3 level free energy profiles of the first and second step of splicing generated while collecting QM/MM+ΔMLP training data. **(a)** Free energy profiles of the first step of splicing with GTP bound with various acids protonating the leaving group. **(b)** Free energy profiles of the first step of splicing comparing G, GTP, and UCG at the $G_{exo}$ position and the impact of Mg(B). **(c)** Free energy profiles of the second step of splicing under varied M(B) and acid coordination conditions.

### Validation of the QM/MM+ΔMLP model

In this work we performed QM/MM simulations with a ΔMLP model trained to correct DFTB3/3ob to the PBE0/6-31G* level within a 6 Å cutoff of the QM region. To validate the model, we first computed the deviation in forces predicted between 4 stochastically trained model replicas, otherwise known as model deviation. The equation for model deviation is given by Eq 7. In the pool-based active learning approach of the DP-GEN simplify program, the commonly accepted threshold for accuracy is to achieve model deviation below 1.84 kcal·mol$^{-1}$, or 0.08 ev/Å.[S9,S10] This is a measure of uncertainty in the prediction made by the model. In the training procedure, an initial batch of 20,000 structures was considered for training. Deviation was computed with the resulting models on the total pool of over 200,000 structures yielding 96.6% of samples with model deviation below the accuracy threshold as shown in Figure S15. In addition, we explicitly compared the predicted versus actual difference in the magnitude of forces between DFTB3 and PBE0 for the training set of 20,000 structures, and a test set of 10,000 non-trained structures. The RMSE values of 0.31 and 0.37 kcal·mol$^{-1}$Å$^{-1}$ respectively indicate high model accuracy across all of the mechanistic scenarios studied, and good transferability beyond the training set.

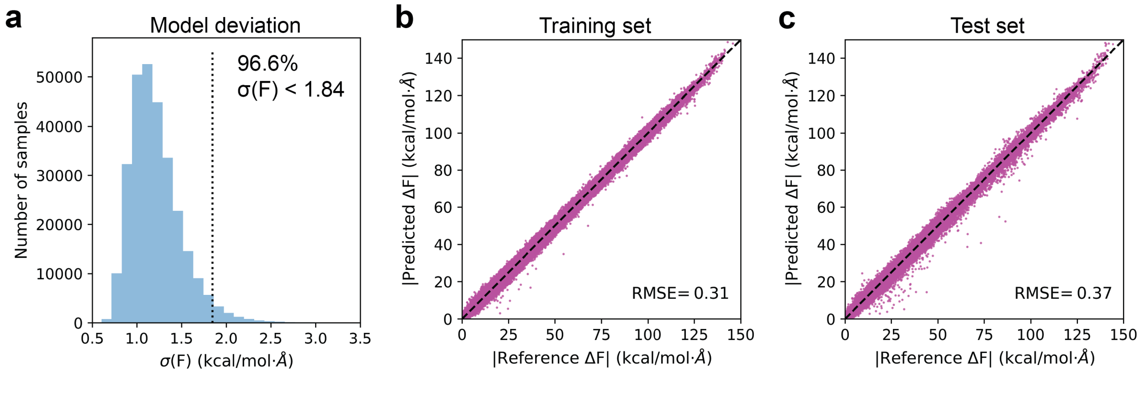


Figure S15. Validation of QM/MM+ΔMLP model for THr reactions. **(a)** Deviation in predicted forces between 4 stochastically trained models. The black vertical line indicates the accuracy cutoff below which samples are removed from the training set and deemed accurate. Comparison of the difference in the magnitude of atomic forces between PBE0 and DFTB3 (reference) and the ΔMLP predicted forces for the **(b)** training **(c)** test set. RMSE is given in kcal·mol$^{-1}$Å$^{-1}$.

### Supplemental QM/MM+ΔMLP level free energy profiles

Here we provide a complete set of free energy profiles obtained for Step 1 with GTP and for Step 2. These include mechanistic scenarios involving acids and bases that were deemed energetically infeasible. Throughout MD simulations, the nucleophillic O3′ in Step 1 forms a hydrogen bond with a water that indirectly coordinates Mg(C). It was not immediately clear whether the high p$K_a$ obtained experimentally arose from the O3′ or this specific water with a stabilizing hydrogen bond and coordination network deprotonating to form hydroxide. Therefore, deprotonation of O3′ by this specific water substituted for hydroxide was included as a reaction coordinate. As discussed in the previous section, DFTB3 level simulations suggested this deprotonation was unfavorable, therefore it was included in the reaction at the QM/MM+ΔMLP level for further investigation. At a higher level of theory, this deprotonation was highly favorable, suggesting the high p$K_a$ is that of O3′. All free energy profiles are given in Figure S16 with barriers provided in Table S7.

In Step 1, the most favorable mechanism involved no general acid, rather Mg(A) acting as a Lewis acid when M(B) was not present was most favorable. This ruled out protonation of the leaving group by a bulk water (Wat), the O2′ of the leaving group sugar which donates a hydrogen bond to O3′, and a water coordinating Mg(A) (Mg(A).Wat). In Step 2, there were less potential acid sources, as Mg(C) does not coordinate a water well positioned to interact with the leaving group, and the O2′ of the $G_\omega$ sugar hydrogen bonds to a nearby phosphate rather than O3′. This left the water indirectly coordinating Mg(C), which has a high p$K_a$ based on Step 1, but could still have a lower p$K_a$ than the leaving group O3′ if its downshift was smaller than that of the nucleophile in Step 1. As shown in Figure S16, it was ultimately unfavorable to deprotonate this water, evidenced by the upturn in the black curve in Figure S16d near the end of the reaction.

One subtle structural difference between Steps 1 and 2 is the orientation of the nucleophile and leaving group sugars relative to solvent, and therefore the ability to hydrogen bond M(B) bound waters. If the pro-$S_P$ in Step 1 (which coordinates Mg(A) and Mg(C)) defines the

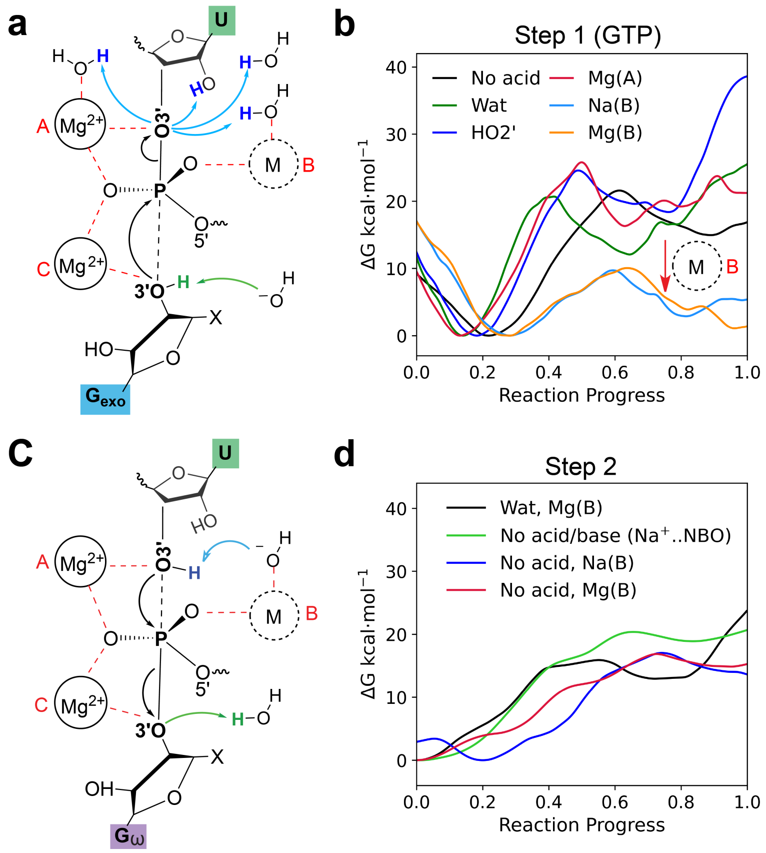


Figure S16. Free energy profiles from ΔMLP accelerated QM/MM simulations of Step 1 and 2 of splicing **(a)** Mechanistic scenarios probed by QM/MM simulations in Step 1. $Mg^{2+}$ ions A and B (marked in red) are always present, while M(B) (dotted circle) is either $Mg^{2+}$, $Na^{+}$, or not present. Phosphoryl transfer is denoted in black, nucleophile deprotonation in green, and leaving group protonation in blue. Metal ion coordination is indicated by red dashed lines. **(b)** QM/MM+ΔMLP level free energy profiles of Step 1 with GTP bound. Legends indicate the acid source being tested, with metals indicating a metal coordinated water. **(c)** Mechanistic scenarios probed in Step 2. **(d)** QM/MM+ΔMLP level free energy profiles of Step 2 with labels corresponding and acid and base sources, and metal indicating a metal coordinated hydroxide.

"ribozyme" side, and pro-$R_P$ the solvent side, the O3′ of $G_{exo}$, and analogously $G_\omega$, is oriented toward the ribozyme, with H2′ facing solvent and sterically blocking solvent side interaction. This orientation is enforced by the simultaneous coordination of Mg(A) by the O3′ and O2′, stabilizing the 3′exo sugar pucker. In Step 2, the O3′ of U is solvent exposed, and can easily hydrogen bond with a water coordinating M(B). This sugar adopts 3′endo, and the O2′ does not coordinate Mg(A). These differences appear to manifest in a few key ways:

Table S7. Free energy barriers of phosphoryl transfer reactions and associated properties for mechanisms studied with QM/MM+ΔMLP simulations

| | | | kcal·mol$^{-1}$ | | Å | kcal·mol$^{-1}$ | | min$^{-1}$ |
|---|---|---|---|---|---|---|---|---|
| System | Acid | Base | $\Delta G^{\ddagger}$ | $\sigma$ | $\xi_1$ | $\Delta G_{\mathrm{rxn}}$ | $\sigma$ | $k_{int}$ |
| Step 1 | | | | | | | | |
| GTP | - | $OH^-$ | 21.59 | 0.17 | 0.99 | 14.98 | 0.31 | $5.47 \times 10^{-2}$ |
| GTP | Wat | $OH^-$ | 20.69 | 0.17 | 0.48 | 24.65 | 0.66 | $3.12 \times 10^{-4}$ |
| GTP | HO2′ | $OH^-$ | 24.57 | 0.50 | 0.44 | 38.50 | 0.28 | $2.20 \times 10^{-14}$ |
| GTP | Mg(A) | $OH^-$ | 25.78 | 0.14 | 0.42 | 21.24 | 0.11 | $4.62 \times 10^{-5}$ |
| GTP,Na(B) | Na(B) | $OH^-$ | 9.72 | 0.10 | 0.43 | 5.28 | 0.17 | $2.77 \times 10^{7}$ |
| GTP,Mg(B) | Mg(B) | $OH^-$ | 10.05 | 0.07 | 0.33 | 1.10 | 0.12 | $1.59 \times 10^{7}$ |
| G | - | - | 16.76 | 0.08 | 0.46 | 15.13 | 0.12 | $1.90 \times 10^{2}$ |
| UCG | - | - | 19.78 | 0.09 | 0.55 | 17.54 | 0.08 | 1.16 |
| UCG | Mg(B).Wat | - | 8.64 | 0.09 | 0.65 | 2.59 | 0.42 | $1.72 \times 10^{8}$ |
| Step 2 | | | | | | | | |
| $G_\omega$,Mg(B) | Wat | Mg(B) | 15.92 | 0.43 | 0.41 | 23.79 | 0.44 | $1.33 \times 10^{-3}$ |
| $G_\omega$ | - | -* | 20.38 | 0.33 | 0.63 | 18.90 | 0.30 | $4.22 \times 10^{1}$ |
| $G_\omega$,Na(B) | - | Na(B) | 17.05 | 0.09 | 0.47 | 13.64 | 0.15 | $1.17 \times 10^{2}$ |
| $G_\omega$,Mg(B) | - | Mg(B) | 16.87 | 0.12 | 0.70 | 14.88 | 0.18 | $1.58 \times 10^{2}$ |

$\Delta G^{\ddagger}$ indicates the rate controlling free energy barrier. Standard error estimates ($\sigma$) were computed from bootstrapping. The reaction coordinate ($\xi_1$) corresponds to the phosphoryl transfer coordinate at the transition state. Positive values indicate the leaving group phosphorous distance exceeds the nucleophile phosphorus distance. Intrinsic reaction rates ($k_{int}$) were estimated from Equation 6 using $\Delta G^{\ddagger}$. A (-) symbol indicates the absence of acid or base. A * indicates a system where $Na^+$ indirectly coordinates the solvent facing NBO at the B site, but does not play an acid or base role. Metals in an acid or base role act through either a water or hydroxide, respectively.

- The reduced inline fitness in Step 2 upon AP formation can be stabilized by a hydrogen bond to Mg(B).Wat, which is sterically blocked in Step 1. Aside from electrostatics, the relative reduction in fitness for Step 2 compared to Step 1 could arise in part from the fact that U forms one stabilizing coordination bond with a metal, while $G_{\mathrm{exo}}$ forms two.

- The free energy profiles of Step 1 have a well defined product minima because the leaving group can be protonated by Mg(B).Wat, or form a stabilizing hydrogen bond on the solvent facing side in addition to HO2′. The product minima in Step 2 is less defined due to lack of solvent exposure, and therefore access to specific or Lewis acid catalysis, as well as lack of stabilization from HO2′.

Given these differences, it could be the case that the product of Step 2 is not protonated directly in the active site, but perhaps upon release.

### Extended analysis of G·U wobble base pair substitution

We performed MD simulations of base pair mutations at the G22·U(-1) site in the first step of splicing to probe whether a synthetic nucleobase could allow accommodation of C at the (-1) position at neutral pH. Taken together, the 3D-RISM, MD, and QM/MM results suggested that when GTP occupies the $G_{exo}$ site, a transient interaction with Mg(B) could occur to trigger catalysis depending on the $Mg^{2+}$ conditions. Given this result, we repeated the MD simulations of the G·C, G·$C^+$, and 7cdiA$P^+$·C containing systems in the presence of Mg(B) to ensure it would not impede the hydrogen bonding network. In fact, the same trend was observed with and without Mg(B), with the G·$C^+$ pair experiencing a break down of the hydrogen bond network that was maintained with the other pairs.

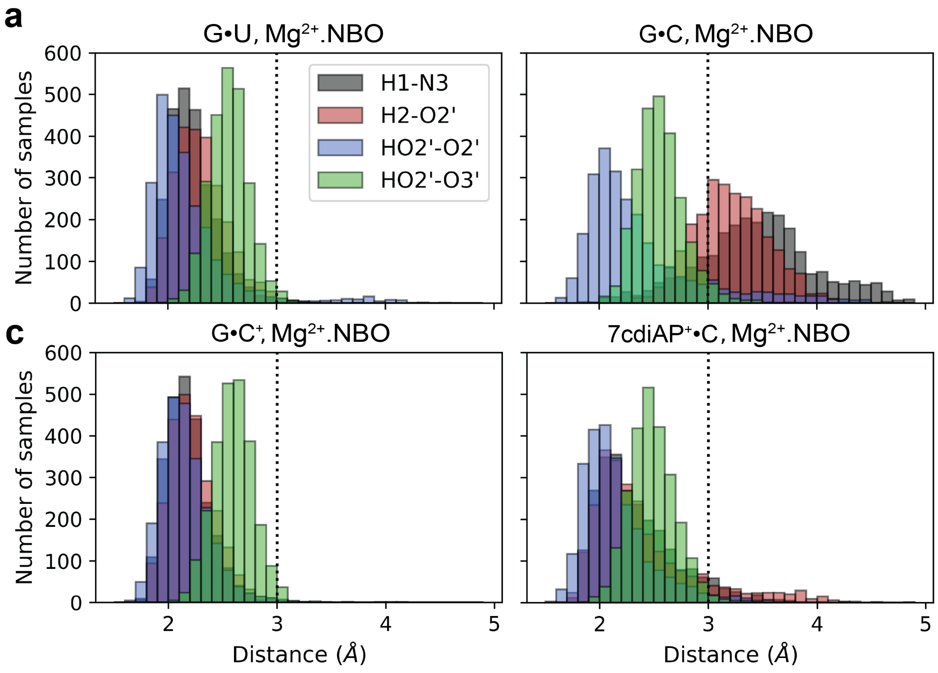


Figure S17. Mutation of the conserved active site G·U wobble base pair upstream of the splice site when Mg(B) is present and coordinating the pro-$R_P$ NBO in the first step of splicing. Histograms quantifying the hydrogen bond distances in the **(a)** G·U, **(b)** G·$C^+$, **(d)** G·C, and **(d)** 7cdiA$P^+$·C containing systems.

### *ab initio* p$K_a$ calculation

To predict the p$K_a$ values for chemically modified diAP variants 3cdiAP and 3,7cdiAP, a series of nitrogen-containing bases (Table S8) with experimentally measured p$K_a$ values was selected as reference and deprotonation free energies were calculated as described in the Methods section and in ref S11. These data were then used to fit a regression model for predicting experimentally unknown p$K_a$ values.

To ensure consistency, reference compounds exhibiting the same protonation pattern—namely, neutral deprotonated structures and positively charged protonated states with experimentally determined p$K_a$ values were selected as benchmarks for p$K_a$ prediction. The results are summarized in Table S8.

A linear regression analysis was performed on these nine data points, yielding the relationship: $y = 0.671x + 2.021$, with $R^2 = 0.9626$, where y represents the experimental p$K_a^{\text{expt}}$ and x represents the calculated p$K_a^{\text{calc}}$. The resulting fitting curve is shown in Figure S18, demonstrating a strong linear correlation between calculated and experimental p$K_a$ values.

To further assess the predictive accuracy of this model, 5 additional compounds (Table S8) with experimentally determined p$K_a$ values exhibiting analogous protonation patterns, specifically, neutral deprotonated and positively charged protonated states were selected as a test set. The same QM computational protocol was applied, and p$K_a$ values were predicted using the established regression equation. All the prediction errors p$K_a^{\text{Error}}$ of the test set, defined as the difference between p$K_a^{\text{model}}$ and p$K_a^{\text{expt}}$ values, were found to be less than 1 p$K_a$ unit, indicating that the model is both reliable and transferable.

With the validated regression model, we performed the same QM computational protocol for diAP and its chemically modified variants 3cdiAP and 3,7cdiAP. Their p$K_a$ results are shown in Table S9.

From a chemical prospective, the p$K_a$ shift at the same site across different structures is expected to be minimal, therefore the average p$K_a^{\text{Error}}$ (-0.30) of diAP and 7cdiAP at N1 position was further considered as a systematic shift of the regression model to predict

Table S8. p$K_a$ values of the fitting set used to construct the regression model and the testing set used for model evaluation

| Dataset | Compound | Position | p$K_a^{calc}$ | p$K_a^{expt}$ | p$K_a^{model}$ | p$K_a^{Error}$ |
|---|---|---|---|---|---|---|
| Fitting | A | N1 | 1.84 | 3.63[S12] | 3.25 | -0.38 |
| | A | N3 | 0.28 | 1.50[S12] | 2.21 | 0.71 |
| | A | N7 | -0.30 | 2.15[S12] | 1.82 | -0.33 |
| | c$^3$A | N1 | 6.74 | 6.80[S13] | 6.55 | -0.25 |
| | c$^7$A | N1 | 4.17 | 5.30[S13] | 4.82 | -0.48 |
| | n$^8$A | N1 | 0.27 | 2.20[S14] | 2.20 | 0.00 |
| | m$^6$A | N1 | 3.29 | 4.01[S15] | 4.23 | 0.22 |
| | m$^{66}$A | N1 | 3.96 | 4.50[S15] | 4.68 | 0.18 |
| | m$^7$G | N1 | 8.21 | 7.20[S16] | 7.53 | 0.33 |
| | | | | | | MAE: 0.32 |
| | | | | | | RMSE: 0.37 |
| Testing | Purine | N1 | 0.81 | 2.40[S17] | 2.56 | 0.16 |
| | 2AP | N1 | 2.73 | 3.80[S18] | 3.85 | 0.05 |
| | C | N3 | 2.61 | 4.20[S19] | 3.77 | -0.43 |
| | 5nC | N3 | -0.48 | 2.64[S20] | 1.70 | -0.94 |
| | 5mC | N3 | 3.36 | 4.30[S21] | 4.28 | -0.02 |
| | | | | | | MAE: 0.32 |
| | | | | | | RMSE: 0.46 |

p$K_a^{calc}$ denotes calculated p$K_a$ values; p$K_a^{expt}$ denotes experimental p$K_a$ values taken from the literature, and p$K_a^{model}$ derives from the regression model. p$K_a^{Error}$ indicates the difference between p$K_a^{model}$ and p$K_a^{expt}$. Adenosine (A), 3-deazaadenosine (c$^3$A), 7-deazaadenosine (c$^7$A), 8-azaadenosine (n$^8$A), 6-methyladenosine (m$^6$A), N6,N6-dimethyladenosine (m$^{66}$A) and N7-methylguanosine (m$^7$G) were selected as fitting set to form the linear regression model ($y = 0.671x + 2.021$, with $R^2 = 0.9626$, where $y$ represents the experimental p$K_a$ values and $x$ represents the calculated p$K_a$ values). Purine, 2-aminopurine (2AP), cytidine (C), 5-azacytidine (5nC) and 5-methylcytidine (5mC) were selected as testing set to validate the regression model. Mean absolute error (MAE) and root mean square error (RMSE) values were reported for both the fitting and testing datasets. Adapted with permission from Puyo-Fourtine, J.; Du, Y.; McCarthy, E.; Ekesan, Ş.; York, D. M. Origins of Reactivity in SAM-Utilizing Ribozyme SAMURI-Catalyzed RNA Alkylation. J. Am. Chem. Soc. 2026, 148 (29), 31316–31330. Copyright 2026 American Chemical Society.

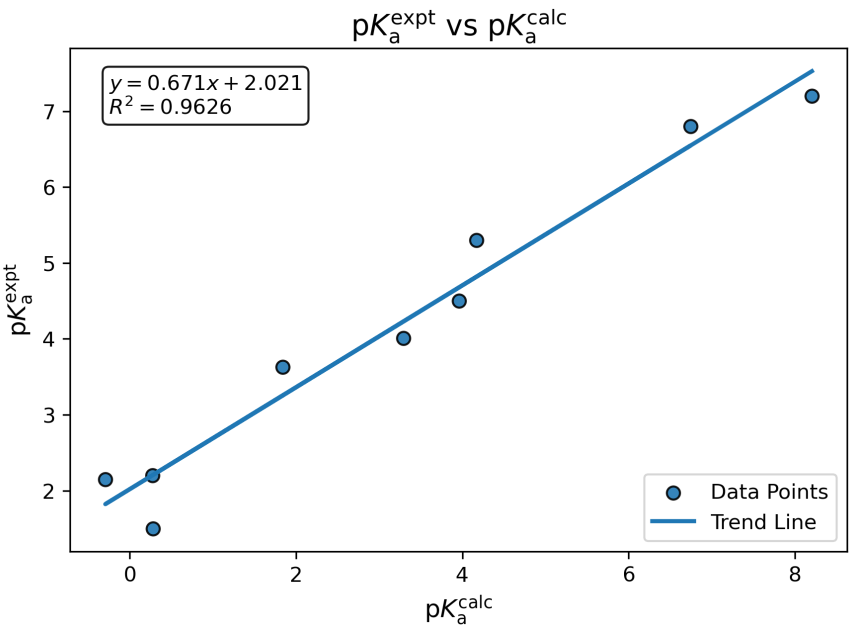


Figure S18. Experimental $\mathrm{p}K_\mathrm{a}$ ($\mathrm{p}K_\mathrm{a}{}^\mathrm{expt}$) vs. calculated $\mathrm{p}K_\mathrm{a}$ ($\mathrm{p}K_\mathrm{a}{}^\mathrm{calc}$). Adapted with permission from Puyo-Fourtine, J.; Du, Y.; McCarthy, E.; Ekesan, Ş.; York, D. M. Origins of Reactivity in SAM-Utilizing Ribozyme SAMURI-Catalyzed RNA Alkylation. J. Am. Chem. Soc. 2026, 148 (29), 31316–31330. Copyright 2026 American Chemical Society.

the $\mathrm{p}K_\mathrm{a}$ values at N1 position. Hereafter, the predicted $\mathrm{p}K_\mathrm{a}$ values are determined by Equation S1 and shown in Figure S19.

$$\begin{aligned}\mathrm{p}K_{\mathrm{a}\,\mathrm{X:N1}}^{\mathrm{pred}} &= \frac{1}{2}\Big[\left(\mathrm{p}K_{\mathrm{a}\,\mathrm{X:N1}}^{\mathrm{model}} - \mathrm{p}K_{\mathrm{a}\,\mathrm{diAP:N1}}^{\mathrm{model}} + \mathrm{p}K_{\mathrm{a}\,\mathrm{diAP:N1}}^{\mathrm{expt}}\right) \\ &\quad + \left(\mathrm{p}K_{\mathrm{a}\,\mathrm{X:N1}}^{\mathrm{model}} - \mathrm{p}K_{\mathrm{a}\,\mathrm{7cdiAP:N1}}^{\mathrm{model}} + \mathrm{p}K_{\mathrm{a}\,\mathrm{7cdiAP:N1}}^{\mathrm{expt}}\right)\Big] \\ &= \mathrm{p}K_{\mathrm{a}\,\mathrm{X:N1}}^{\mathrm{model}} + 0.30.\end{aligned} \tag{S1}$$

Table S9. $\mathrm{p}K_\mathrm{a}$ values and prediction errors for isofunctional nucleotides of interest

| Compound | Position | $\mathrm{p}K_\mathrm{a}^\mathrm{calc}$ | $\mathrm{p}K_\mathrm{a}^\mathrm{expt}$ | $\mathrm{p}K_\mathrm{a}^\mathrm{model}$ | $\mathrm{p}K_\mathrm{a}^\mathrm{Error}$ | $\mathrm{p}K_\mathrm{a}^\mathrm{pred}$ |
|---|---|---|---|---|---|---|
| diAP | N1 | 4.22 | 5.10[S18] | 4.85 | -0.25 | - |
| 7cdiAP | N1 | 4.96 | 5.70[S22] | 5.35 | -0.35 | - |
| 3cdiAP | N1 | 6.48 | - | 6.37 | - | 6.67 |
| 3,7cdiAP | N1 | 7.65 | - | 7.16 | - | 7.46 |

$\mathrm{p}K_\mathrm{a}^\mathrm{calc}$ denotes calculated $\mathrm{p}K_\mathrm{a}$ values; $\mathrm{p}K_\mathrm{a}^\mathrm{expt}$ denotes experimental $\mathrm{p}K_\mathrm{a}$ values taken from the literature, and $\mathrm{p}K_\mathrm{a}^\mathrm{model}$ derives from the regression model. $\mathrm{p}K_\mathrm{a}^\mathrm{Error}$ indicates the difference between $\mathrm{p}K_\mathrm{a}^\mathrm{model}$ and $\mathrm{p}K_\mathrm{a}^\mathrm{expt}$. $\mathrm{p}K_\mathrm{a}^\mathrm{pred}$ denotes the final predicted values.

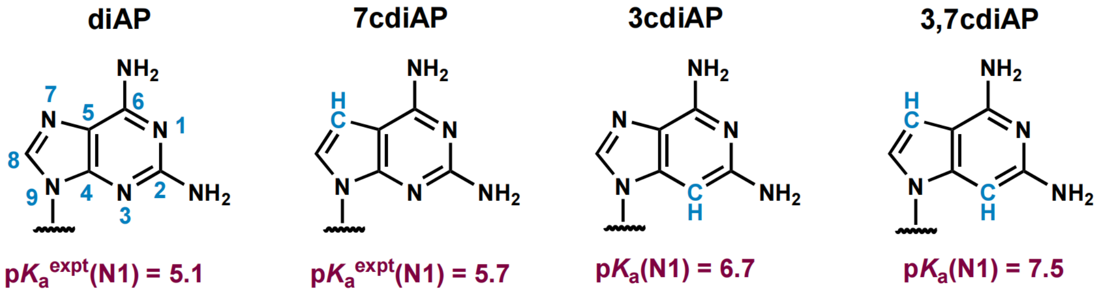


Figure S19. p$K_a$s predicted from *ab initio* quantum chemical calculations for potential mutants of G22 to support wobble base pairing with C(-1).

### Details of system builds

Table S10. Details of all systems builds including non-standard residues and solvent components

| Prev seq | Nuc | 22·(-1) pair | B site w/o Mg | | | B site w Mg | | |
|---|---|---|---|---|---|---|---|---|
| | | | Na | Cl | Mg | Na | Cl | Mg |
| Step 1, 7YCH | | | | | | | | |
| 1: 405 nucleotides, 69 906 waters | | | | | | | | |
| | GTP | G·U | 524 | 180 | 31 | 522 | 180 | 32 |
| | $GTP^-$ | G·U | 524 | 179 | 31 | 522 | 179 | 32 |
| | $GTP^-$ | G·C | 524 | 179 | 31 | 522 | 179 | 32 |
| | $GTP^-$ | $G·C^+$ | 523 | 179 | 31 | 521 | 179 | 32 |
| | $GTP^-$ | $7cdiAP·C^+$ | 523 | 179 | 31 | 521 | 179 | 32 |
| | $G^-$ | G·U | 520 | 179 | 31 | not built | | |
| 2: 407 nucleotides, 65 156 waters | | | | | | | | |
| UC | G | G·U | 511 | 169 | 31 | 509 | 169 | 32 |
| UC | $G^-$ | G·U | 511 | 168 | 31 | 509 | 168 | 32 |
| Step 2, 7YGA | | | | | | | | |
| 3: 407 nucleotides, 66 956 waters | | | | | | | | |
| 5′ exon | U | G·U | 514 | 172 | 31 | 512 | 172 | 32 |
| 5′ exon | $U^-$ | G·U | 514 | 171 | 31 | 512 | 171 | 32 |

Non-standard residues with a (-) charge are deprotonated at the O3′ position, $C^+$ is protonated at the N3 position, and $7cdiAP^+$ is protontated at the N1 position.

### Convergence of MD simulations: root mean square deviation analysis

In order to characterize the dynamical ensemble of the active, reactant state in the first and second steps of splicing under various conditions we conducted 5 independent MD simulations for each mechanistically distinct state. Convergence of these simulations was assessed by analyzing the root mean square deviation (RMSD) of the heavy atoms' positions with respect to the first frame of the trajectory. RMSD analysis over time indicates that the simulations generally reached convergence after ∼100 ns, therefore the first 100 ns were discarded as equilibration and further analysis was done on the final 100 ns. The RMSD from simulations of the first step of splicing are given in Figure S20. In general, RMSD values in the production region ranged from 2 to 6 Å. When $UCG_{exo}$ is present, the RMSD values are generally lower than for G or GTP, likely due to stabilizing base pairing with the ribozyme.

The same set of MD simulations was repeated for the second step, except the $G_{exo}$ position is now occupied by $G_{\omega}$ of the UCG substrate, and was therefore not varied. In order to equilibrate the standard state of step 2, an initial 100 ns of simulations with restraints of 2.5 Å between Mg(C) and its coordinating phosphates was required to maintain the ion in its binding site. Restraint weights were held at 25 $kcal{\cdot}mol^{-1}Å^{-1}$ for 25 ns, and reduced to 0 over the next 75 ns. This ion was not resolved in the cryo-EM due to a clashing phosphorothioate substitution, therefore the initial positions yielded close contacts with the ion that required gradual equilibration. This restrained simulation time was not considered for RMSD analysis, and the next 100 ns of unrestrained simulation was discarded as equilibration.

In addition, three different combinations of base pairing were tested at the G22·U(-1) wobble position in the first step of splicing. These departed from the GTP bound AP state either with or without Mg(B). The associated RMSD values are shown in Figure S22. The presence of Mg(B) seemingly had a stabilizing effect for the G·C and $7cdiAP^{+}{\cdot}C$ base pairings, but less so for $G{\cdot}C^{+}$. The (-1) residue is solvent exposed, and therefore closer to the B site. A positive charge at this position may influence potential Mg(B) binding, less so than nucleotide 22 which is shielded from solvent.

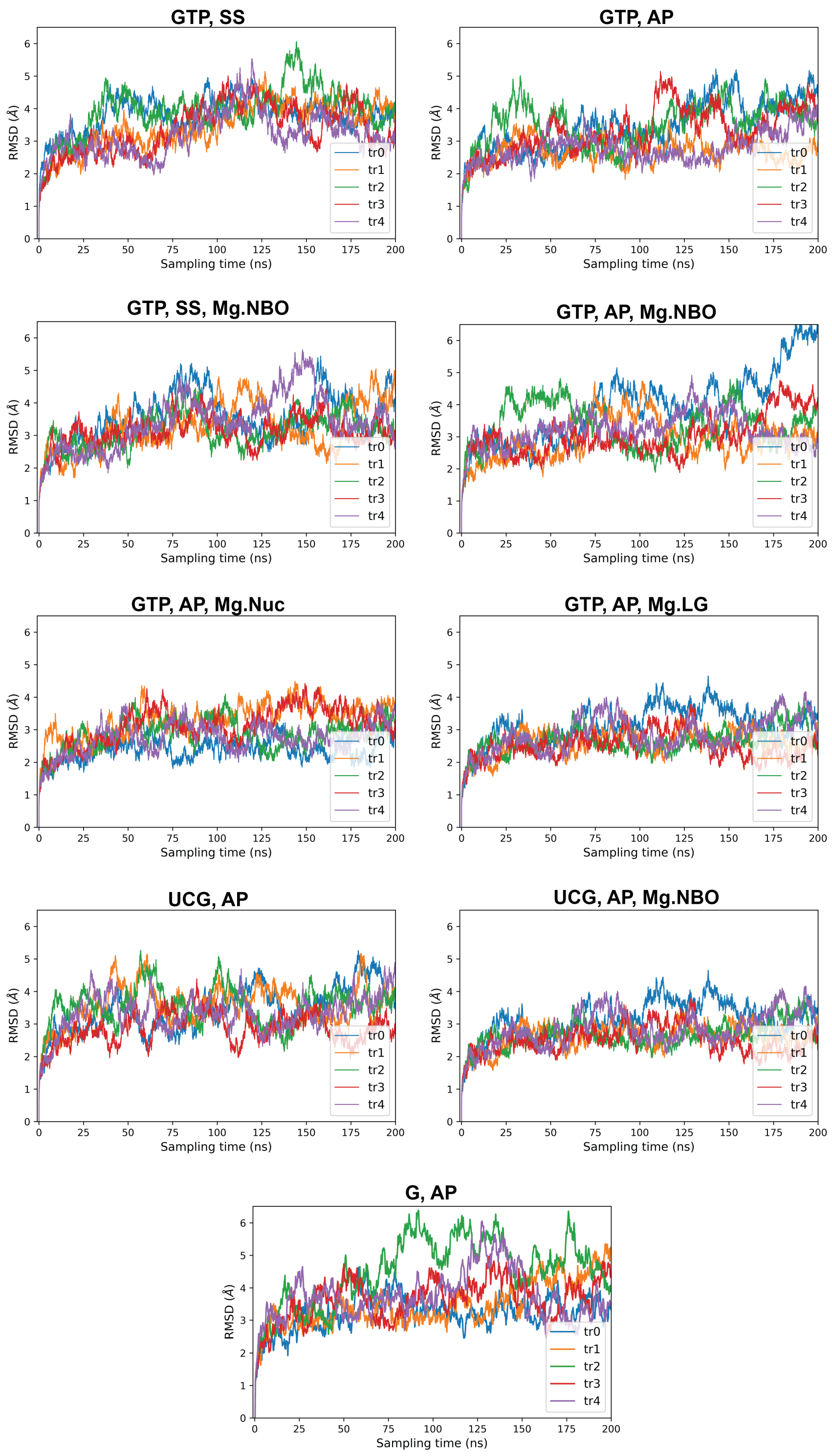


Figure S20. RMSD analysis of heavy atom positions relative to the first frame of 5 independent 200 ns MD trials for the first step of splicing. Titles indicate the $G_{exo}$ moiety, protonation state of the nucleophile, and Mg(B) binding mode if present. SS indicates the standard state with nucleophile protonated, and AP indicates the activated precursor with nucleophile deprotonated. Mg.X indicates direct coordination of Mg(B) to the species X.

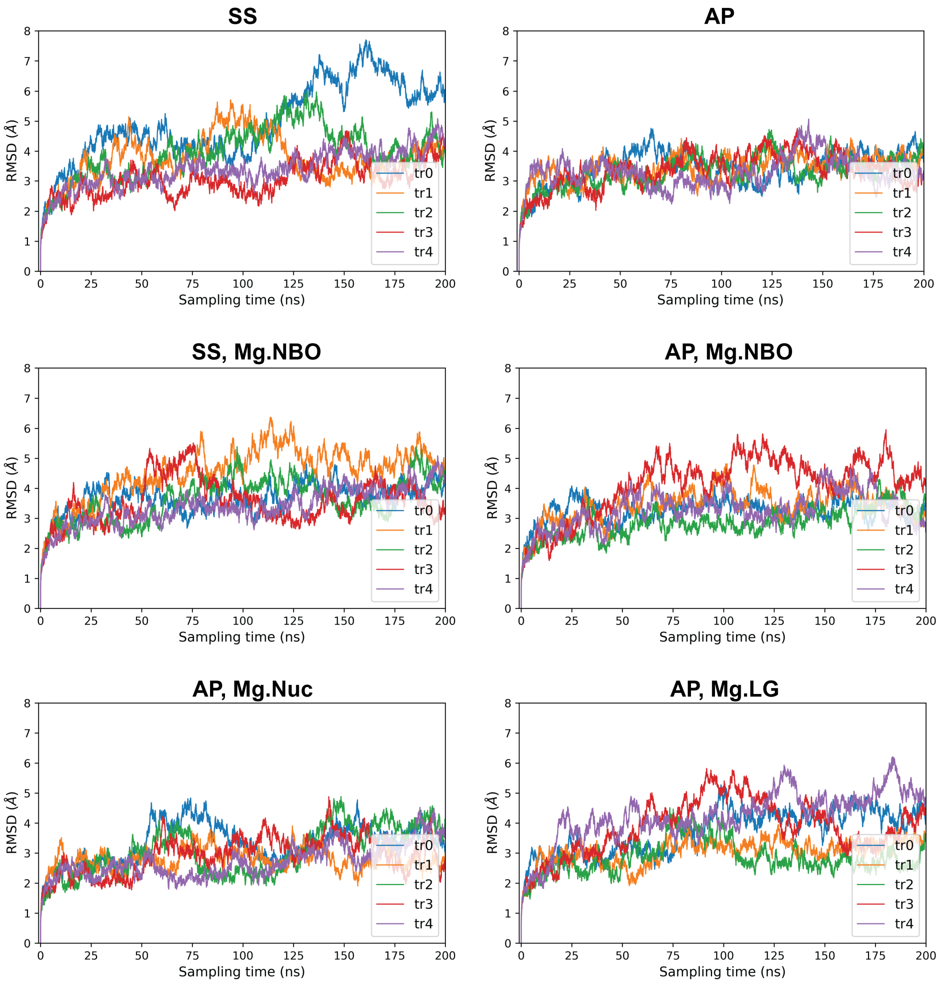


Figure S21. RMSD analysis of heavy atom positions relative to the first frame of 5 independent 200 ns MD trials for the second step of splicing. Titles indicate protonation state of the nucleophile and Mg(B) binding mode if present.

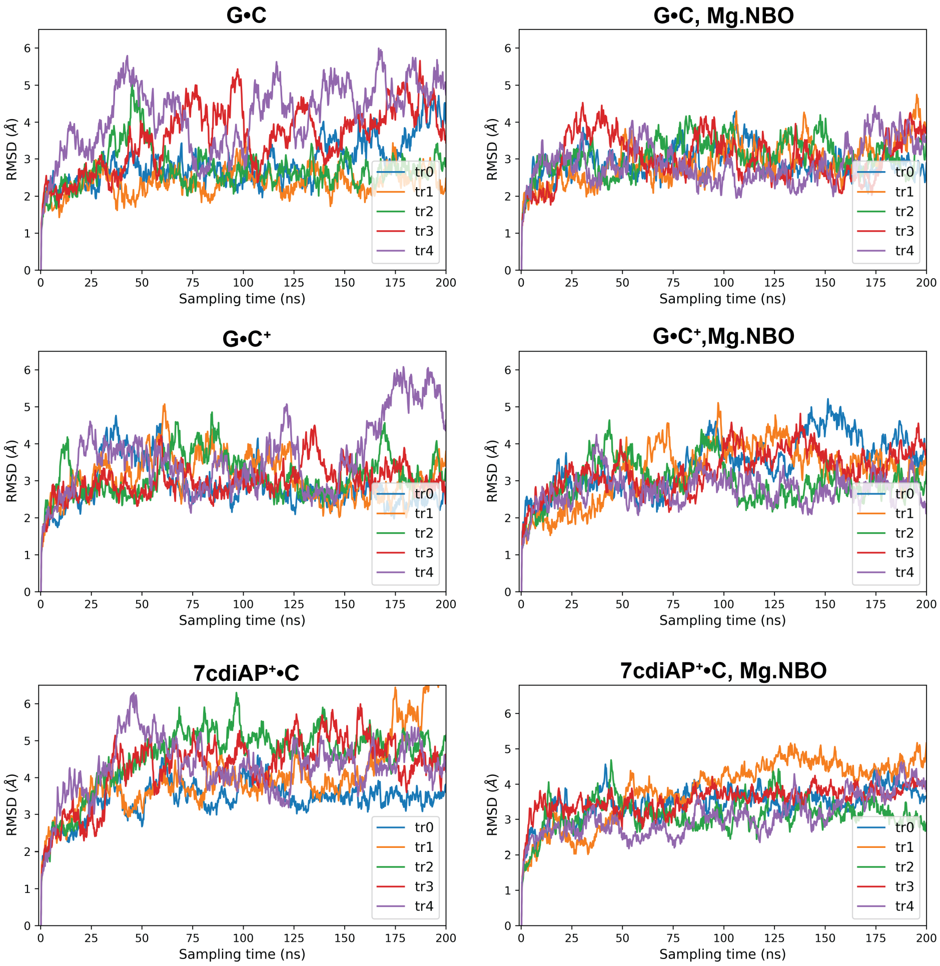


Figure S22. RMSD analysis of heavy atom positions relative to the first frame of 5 independent 200 ns MD trials for the first step of splicing with G·U wobble base pair substitutions. Titles indicate base pairing at the (-1) position, and Mg(B) binding mode if present. All simulations represent the AP state with GTP bound.

(S15) Martin, D. M. G.; Reese, C. B. Some aspects of the chemistry of N(1)- and N(6)-

dimethylallyl derivatives of adenosine and adenine. *J. Chem. Soc. C* **1968**, *0*, 1731–1738.

(S16) Lawley, P. D.; Brookes, P. Further Studies on the Alkylation of Nucleic Acids and their Constituent Nucleotides. *Biochem. J.* **1963**, *89*, 127–38.

(S17) Kampf, G.; Kapinos, L. E.; Griesser, R.; Lippert, B.; Sigel, H. Comparison of the acidbase properties of purine derivatives in aqueous solution. Determination of intrinsic proton affinities of various basic sites. *J. Chem. Soc. Perkin Trans. 2* **2002**, *2*, 1320–1327.

(S18) Han, J.; Burke, J. M. Model for General Acid-Base Catalysis by the Hammerhead Ribozyme: pH-Activity Relationships of G8 and G12 Variants at the Putative Active Site. *Biochemistry* **2005**, *44*, 7864–7870.

(S19) Shugar, D.; Fox, J. J. Spectrophotometric studies of nucleic acid derivatives and related compounds as a function of pH. *Biochim. Biophys. Acta.* **1952**, *9*, 199–218.

(S20) Notari, R. E.; DeYoung, J. L. Kinetics and Mechanisms of Degradation of the Antileukemic Agent 5-Azacytidine in Aqueous Solutions. *J. Pharm. Sci.* **1975**, *64*, 1148–57.

(S21) Fox, J. J.; Van Praag, D.; Wempen, I.; Doerr, I. L.; Cheong, L.; Knoll, J. E.; Eidinoff, M. L.; Bendich, A.; Brown, G. B. Thiation of Nucleosides. II. Synthesis of 5-Methyl-2'-deoxycytidine and Related Pyrimidine Nucleosides¡sup¿1¡/sup¿. *J. Am. Chem. Soc.* **1959**, *81*, 178–187.

(S22) Seela, F.; Peng, X. 7-Functionalized 7-Deazapurine Ribonucleosides Related to 2-Aminoadenosine, Guanosine, and Xanthosine: Glycosylation of Pyrrolo[2,3-*d*]pyrimidines with 1-*O*-Acetyl-2,3,5-tri-*O*-benzoyl-D-ribofuranose. *J. Org. Chem.* **2006**, *71*, 81–90.
